# Müller Glia regenerative potential is maintained throughout life despite neurodegeneration and gliosis in the ageing zebrafish retina

**DOI:** 10.1101/2020.06.28.174821

**Authors:** Raquel R. Martins, Mazen Zamzam, Mariya Moosajee, Ryan Thummel, Catarina M. Henriques, Ryan B. MacDonald

## Abstract

Ageing is a significant risk factor for degeneration of the retina. Harnessing the regenerative potential of Müller glia cells (MG) in the retina offers great promise for the treatment of blinding conditions. Yet, the impact of ageing on MG regenerative capacity has not yet been considered. Here we show that the zebrafish retina undergoes telomerase-independent age-related neurodegeneration. Yet, this progressive neuronal loss in the ageing retina is insufficient to stimulate the MG regenerative response. Instead, age-related neurodegeneration leads to MG gliosis and loss of vision, similarly to humans. Nevertheless, gliotic MG cells retain Yap expression and the ability to regenerate neurons after acute light damage. Therefore, we identify key differences in the MG response to acute versus chronic damage in the zebrafish retina and show that aged gliotic MG can be stimulated to repair damaged neurons in old age.

**SUMMARY:** 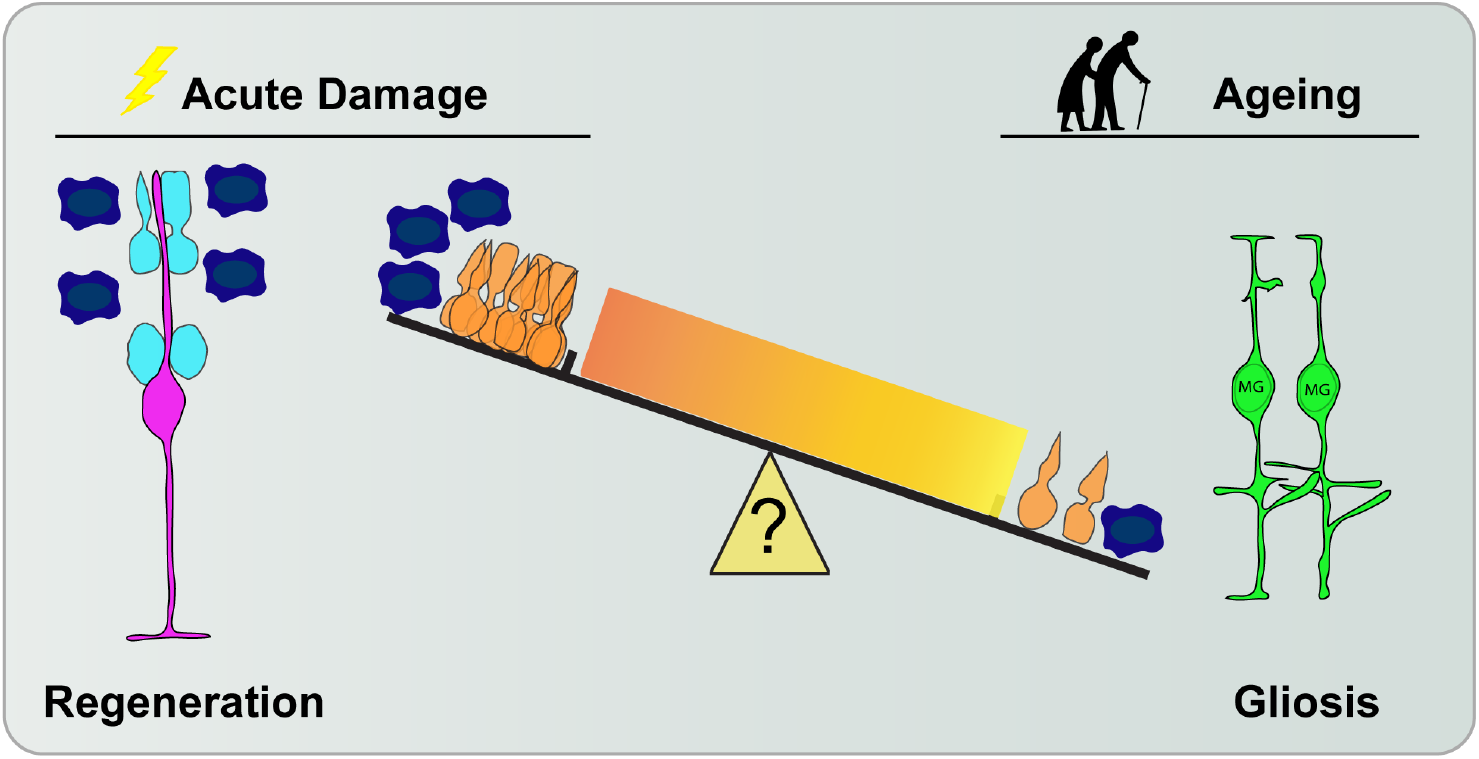

Our data suggest there are key differences between mechanisms driving regeneration in response to acute damage versus age-related chronic damage. It may be that either the number of cells dying in natural ageing is not enough to stimulate MG to proliferate, or the low number of microglia and respective signals released are not sufficient to trigger MG proliferation. Importantly, we show that gliotic MG cells can be stimulated to repair damaged neurons in old zebrafish retina.

## INTRODUCTION

The physiology and structure of the healthy human eye is well known to degrade with age. As the population is ageing worldwide, the identification and potential treatment of the underlying cellular and molecular dysfunctions in the aged retina will be critical for global health going forward ^1^. Hallmarks of the ageing retina include tissue thinning, neuronal loss and reduced visual function, especially in the macula ^2–7^. Further, ageing is a significant risk factor for degeneration and disease in the retina, such as age-related macular degeneration or primary open angle glaucoma^6–8^. Retinal degeneration is often a process that plays out over months or years, whereby neurons gradually die leading to dysfunction ^5,6^. Regenerative-based therapies, such as stimulating endogenous Müller Glia (MG) to regenerate, offer great promise for the treatment of various types of blindness resulting from the loss of retinal neurons. Thus, determining the effects of ageing on the regenerative potential of the retina is an important consideration for the efficacy of such treatments going forward.

The principal cells tasked with maintaining the retina throughout life are the MG cells. MG provide retinal neurons with a myriad of support functions, including trophic support, neurotransmitter recycling and energy metabolism ^9^. While these functions are critical for healthy retinal function, MG also have a prominent role during disease and after neuronal insult. In the mammalian retina, neuronal damage results in a MG gliotic response, involving the up-regulation of stress proteins, proliferation and morphology changes. This response is thought to be neuroprotective initially, but may ultimately culminate in dysfunction and death via loss of metabolic support or tissue integrity ^10^. However, in “lower vertebrates”, such as fish and amphibians, the MG cells have the capacity to regenerate the retina after neuronal damage ^11^. Intensive efforts have been made to uncover the molecular mechanisms regulating these endogenous regenerative responses in the retinas of lower vertebrates ^12–17^. In the zebrafish retina, after acute damage, MG undergo a brief reactive gliosis-like phase that transitions into a regenerative response: a de-differentiation and proliferation of the MG progenitor cell to specifically generate new neurons and restore vision ^18–20^. While the mammalian retina lacks significant regenerative capacity ^21,22^, re-introduction of key molecular signals into MG has shown great promise to stimulate the genesis of new neurons in mice after damage ^23–25^. These molecular mechanisms are largely studied in the young adult zebrafish using acute injury models, such as those generated by intense light or toxins ^14,26^. In contrast to acute injury, damage to the retina caused by degenerative disease or in ageing manifest themselves over much longer timescales and result in the gradual accumulation of DNA damage and cell death^7^. As such, chronic degeneration may significantly differ from injury models, especially in its capacity to induce a regenerative response from glia. While multiple acute injuries do not appear to reduce the overall regenerative response of MG^26^, the effects of natural chronic age-related damage on the MG regenerative potential has not been determined.

The zebrafish is an emerging ageing model, as it displays key human-like hallmarks of ageing in most tissues, including the retina ^27,28^. These include an age-related decrease in cell density and increased cell death, suggesting chronic degeneration of the retina with increasing age ^27^. Depending on the tissue, hallmarks of ageing in the zebrafish include telomere shortening, DNA damage, decreased proliferation and apoptosis or senescence ^29–31^. Accordingly, the telomerase mutant (*tert^-/-^*) zebrafish model is now an established accelerated model of ageing ^29–31^, where regeneration is known to be impaired in tissues such as the heart ^32^. The *tert^-/-^* model therefore offers the possibility of studying key aspects of ageing in a shorter time period, and allows for the identification of telomerase-dependent mechanisms of ageing, potentially involved in retinal regeneration ^29–31^.

Here, we used the zebrafish retina as a model study the impact of ageing on the maintenance of neuronal structure and regenerative capacity of MG cells. We show that naturally aged and telomerase deficient retinas display known hallmarks of retinal ageing, including tissue thinning, accumulation of DNA damage and neuronal death. However, this accumulation of chronic damage does not stimulate MG to proliferate or replace lost neurons. Instead, and in contrast to acute injury, ageing leads to a gliotic response in MG and loss of vision, recapitulating hallmarks of human retinal degeneration with advancing age. Yet, gliotic MG in the old retinas retain the expression of Hippo signalling, and thereby their potential for regeneration. After acute light damage, gliotic MG cells maintain their regenerative potential to old ages, and are therefore are capable of regenerating neurons throughout life. We therefore identify key differences in the MG regenerative response to acute versus chronic damage, a key consideration in potential therapies aiming to stimulate endogenous regenerative mechanisms to treat human retinal disease.

## RESULTS

### The aged zebrafish retina displays neurodegeneration independently of telomerase

Similar to humans, the zebrafish retina consists of three nuclear layers separated by two synaptic plexiform layers. The nuclear layers consist in the outer nuclear layer (ONL), containing photoreceptors; the inner nuclear layer (INL) containing bipolar cells (BCs), amacrine cells (ACs), horizontal cells (HCs) and MG; and the ganglion cell layer (GCL) mainly containing retinal ganglion cells (RGCs) (see diagram in Figure 1A). The zebrafish retina is considered specified and functional by 73 hours post-fertilisation (hpf) ^33^. The retina continually grows at the periphery due to proliferation of the ciliary marginal zone (CMZ) ^17^, a population of progenitors thought to contribute to retinal growth throughout life. Thus, the central region of the adult retina can be considered the “oldest”, as it is largely generated during early developmental stages, and where there is very little proliferation to generate new neurons ^28^. In contrast, the peripheral region of the retina is continually expanding at the margins due with the newest born cells being found adjacent to the CMZ.

**Figure 1.**
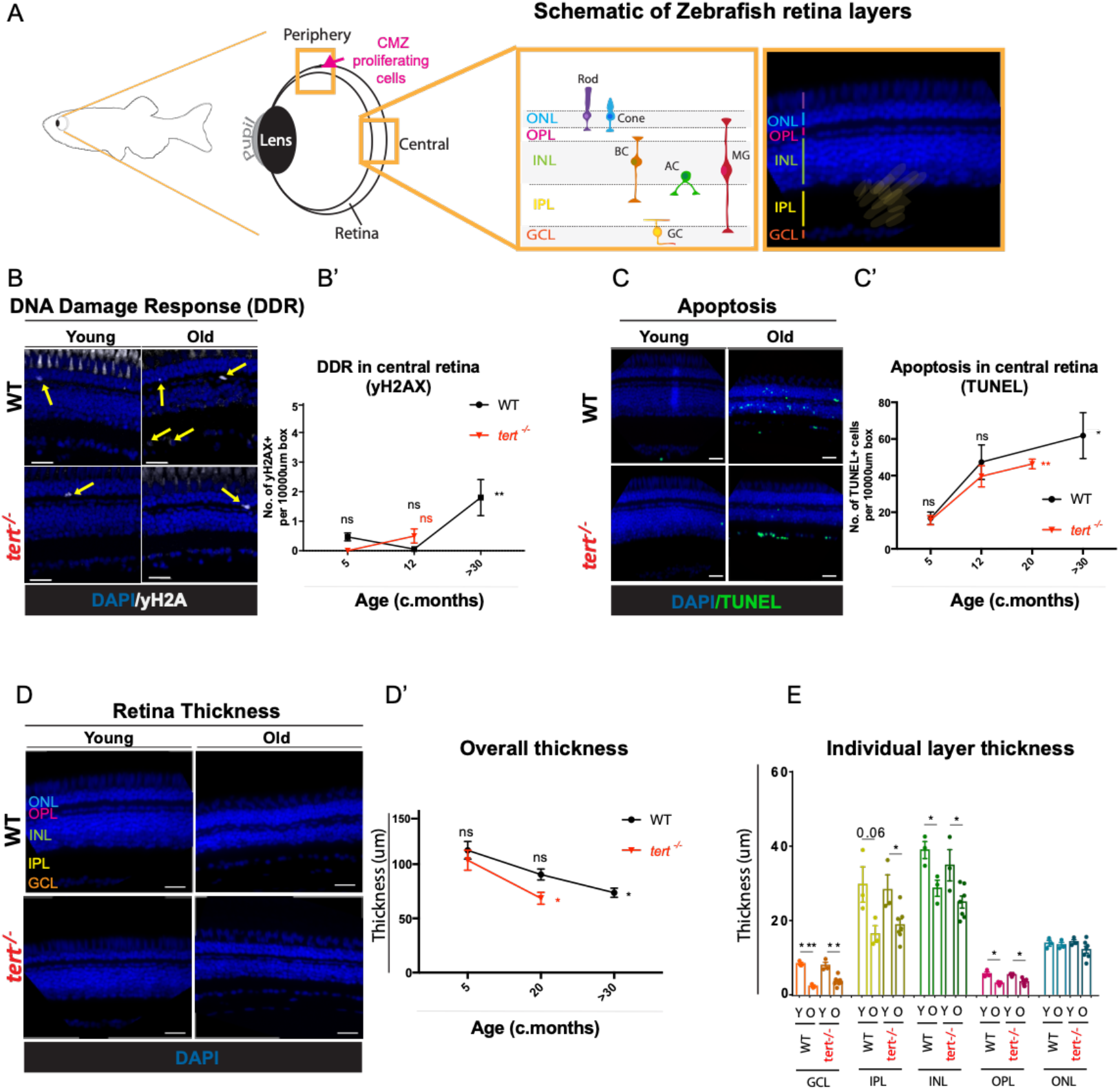
Zebrafish retina ageing is characterised by increased DNA damage, cell death, and tissue thinning over-time, independently of telomerase. (A) Schematic figure of the zebrafish retina highlighting the periphery, including Ciliary Marginal Zone (CMZ) and central retina, with respective layers and cell types (inset). (B-C) The central retina immunolabeled with (B) γH2AX (DNA damage, in white) and (C) TUNEL (apoptotic cells, in green) in both WT and *tert^-/-^*, at young (5 months) and old ages (>30 months in WT and 12 or 20 in *tert^-/-^*). Scale bars: 20μm. (B) Yellow arrows highlight γH2AX^+^ cells. (B’-C’) Quantifications of the number of (B’) γH2AX^+^ cells and (C’) TUNEL^+^ cells per area (10 000μm^2^). Error bars represent the SEM. N=3-4. (D) Central retina thinning in both WT and *tert^-/-^*, at young (5 months) and old ages (>30 months in WT and 20 months in *tert^-/-^*). Scale bars: 20μm. (D’, D’’) Quantifications of the central retina thickness cells per area (10 000μm^2^), (D’) in the overall retina and (D’’) per layer of the retina. Error bars represent the standard error of the mean (SEM). N=3. P-value: * <0.05; ** <0.01; *** <0.001.

It has recently been shown that the ageing zebrafish retina shows some degeneration with advancing age ^27,28^. Telomerase is known to be important for proliferation ^30,31^ and regeneration in the zebrafish ^32^.^32^; however, its specific role in the ageing retina remains unclear. Here, we characterised key hallmarks of ageing, (i.e DNA damage response and apoptosis) in the central and the peripheral retina (Fig 1; Supplementary Fig 1) throughout the zebrafish lifespan in naturally aged wildtype (WT) and telomerase mutants (*tert^-/-^*). The naturally aged WT central retina displays accumulation of DNA damage, as evidenced by the presence of strong γH2AX nuclear foci, a molecular marker of the DNA damage response (Fig 1B, B’). This is accompanied by a significant increase in cell death, as assessed by the early cell death labelling terminal deoxynucleotidyl transferase dUTP nick end labelling (TUNEL) (Fig 1C, C’). Increased DNA damage and cell death with ageing occur in a telomerase-independent manner, as no differences are identified between genotypes at any time-point analysed. Further, retinal thinning is a likely consequence of increased cell death and is a known hallmark of human retinal ageing ^34^. Accordingly, we show that zebrafish central retina progressively thins with ageing, consistent with previous studies ^27,28^. Moreover, as for increased DNA damage and cell death, we show that this occurs largely independently of telomerase (Fig 1D).

We next sought to determine whether this cell death and retinal thinning is caused by the specific loss of any particular neuronal type. We used immunohistochemistry with several molecular markers specific for different neuronal populations throughout the lifespan of zebrafish (Table 1), in the presence and absence of telomerase (*tert^-/-^*). As expected, there is an overall reduction in neuron numbers with ageing in the zebrafish retina (Fig 2A-C). Importantly, this neuronal loss is not due to a decreased number of RGCs in the GCL (Fig 2D), but due to a decreased number of HuC/D-expressing ACs (Fig 2E) and PKC-expressing BCs in the INL layer (Fig 2F). Additionally, in the aged zebrafish retina, BCs display disorganised axon terminals in the IPL (Fig 2A, B). None of these neurodegenerative phenotypes are affected by depletion of telomerase, as there are no differences between WT and *tert^-/-^* at any of the tested ages. The RGCs, ACs and BCs are neurons that come together to make connections in the major synaptic neuropil, called the inner plexiform layer (IPL; Fig 1A). Thus, degeneration observed in these neuronal populations is expected to affect connectivity in the IPL. Accordingly, we confirmed thinning of the synaptic IPL layer due to a reduction and disorganisation of Ribeye A positive pre-synaptic terminals with ageing (Fig 2G). The loss of photoreceptor integrity is one of the key features of human retina ageing and disease ^35^ and we show that rod photoreceptor outer segments in the zebrafish retina undergo dramatic structural changes with ageing (Fig 2H), confirming previous results ^27^. These structural changes are accompanied by disruption of the tight junction protein zonula occludens (ZO-1) (Fig 2I), a marker for the outer limiting membrane thought to be involved in photoreceptor degeneration ^28^. These defects are also not accelerated in the *tert^-/-^* at any age tested, suggesting this is likely a telomerase-independent phenomenon.

**Figure 2:**
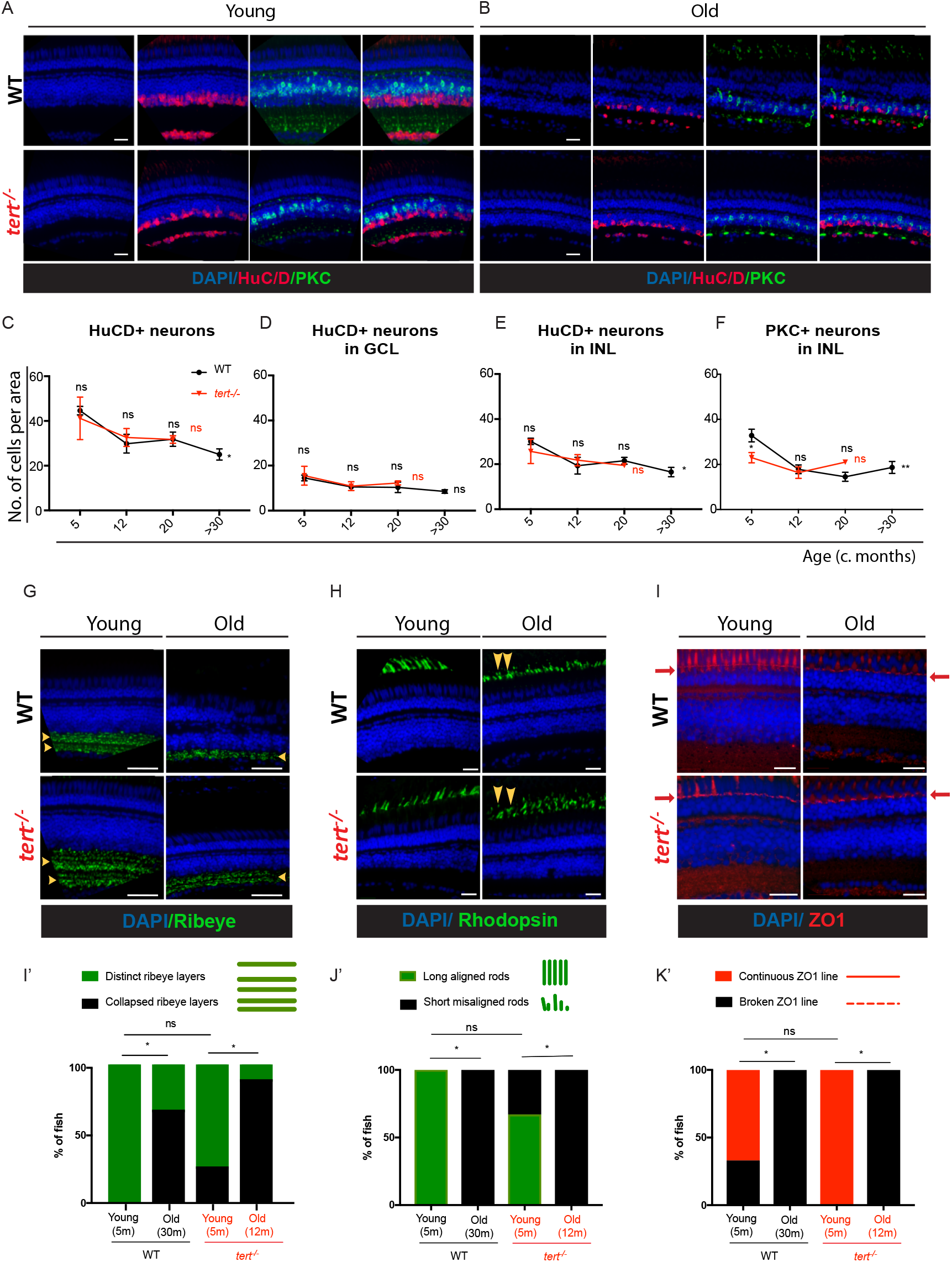
The zebrafish retina undergoes neurodegeneration with ageing, independently of telomerase. The central retina immunolabeled with HuC/D and PKC (amacrine in magenta and bipolar cells in green, respectively), in both WT and *tert^-/-^*, at (A) young (5 months) and (B) old ages (>30 months in WT and 12 months in *tert^-/-^*. Scale bars: 20μm. (C-F) Quantifications of the number of HuC/D-positive neurons per area (10 000μm^2^) (C) in the overall retina, (D) in the GCL (ganglion cells) and (E) in the INL (amacrine cells). (F) Quantifications of the number of PKC-positive neurons cells per area (10 000μm^2^) in the INL (bipolar cells). Error bars represent the SEM. N=3-4. (G-I) The central retina immunolabeled with (G) ribeyeA (pre-synaptic ribbons, in green), (H) rhodopsin (photoreceptors outer segments, in green), and (I) ZO1 (outer limiting membrane, in red), in both WT and *tert^-/-^*, at young (5 months) and old ages (>30 months in WT and 12 months in *tert^-/-^*). Scale bars: 20μm. (G’-I’) Quantification of the percentage of fish presenting defects in (G’) ribeye (collapsed ribeye layers), (H’) rhodopsin (short and misaligned outer segments), and (I’) ZO1 (broken outer limiting membrane). Error bars represent the SEM. (G’) N=4-9, (H’, I’) N=3-6. P-value: * <0.05; ** <0.01; *** <0.001.

**Table 1.** Primary antibodies used for immunostaining.

| **Antibody,**<br>**species and type** | **Dilution**<br>**factor** | **Catalogue number;**<br>**Company, City, Country** |
| --- | --- | --- |
| yH2AX rabbit polyclonal | 1:500 | GTX127342; GeneTex, Irvine, CA, USA |
| p53 rabbit polyclonal | 1:200 | AS-55342s; Labscoop, Little Rock, AR, USA |
| PCNA mouse monoclonal | 1:500 | NB500-106; Novus Biologicals, Littleton, CO, USA |
| PCNA rabbit polyclonal | 1:50 | GTX124496; GeneTex, Irvine, CA, USA |
| 7.4.C4 (4C4) mouse monoclonal | 1:100 | A gift from A. McGown |
| HuCD mouse monoclonal | 1:100 | A21271; Thermo Fisher Scientific, Waltham, MA, USA |
| PKCβ1 rabbit polyclonal | 1:100 | Sc-209; Santa Cruz, Dallas, TX, USA |
| Ribeye rabbit polyclonal | 1:10,000 | A gift from Teresa Nicholson |
| GFAP rabbit polyclonal | 1:200 | Z0334, Agilent DAKO, Santa Clara, CA, USA |
| GFAP mouse monoclonal | 1:100 | zrf1, ZIRC |
| Glutamine Synthase (GS) mouse monoclonal | 1:150 | mab302, Merck, Kenilworth, NJ, USA |
| Rhodopsin rabbit polyclonal | 1:5,000 | A gift from David Hyde |
| BrdU rat | 1:200 | OBT0030A, Accurate Chemical & Scientific, Westbury, NY, USA |
| Yap mouse monoclonal | 1:200 | Sc-101199; Santa Cruz, Dallas, TX, USA |
| Cralbp rabbit polyclonal | 1:200 | 15356-1-AP; Proteintech, Rosemont, Illinois, USA |
| Vimentin mouse monoclonal | 1:500 | 60330-1-IG; Proteintech, Rosemont, Illinois, USA |
| Pax6 rabbit monoclonal | 1:200 | 12323-1-AP Proteintech, Rosemont, Illinois, USA |
| Tgfb1 rabbit polyclonal | 1:100 | Ab215715, Abcam plc. Cambridge, UK |
| Tgfb2 rabbit polyclonal | 1:100 | Ab36495, Abcam plc. Cambridge, UK |
| Tgfb3 rabbit polyclonal | 1:100 | Ab215715, Abcam plc. Cambridge, UK |

### MG do not proliferate in response to chronic retinal neurodegeneration with ageing

It is well established that MG in the zebrafish retina proliferate and regenerate lost neurons after acute injury ^11–13^. Although the progressive age-related retinal neurodegeneration would suggest a lack of regenerative response by MG, it remains unclear whether these degenerations are being counteracted, at least to some degree, by proliferation in the aged retina. To test this, we used an EDU pulse-chase strategy to identify any cell divisions in the retina throughout lifespan, which would be indicative of potential regeneration by MG. To characterise the steady state regenerative capacity of the central and peripheral CMZ until old age, we carried out a 3-day pulse of EdU followed by 0- and 30-days chase, in young and old WT and *tert^-/-^* zebrafish (Fig 3A, B). We observe few EdU-positive cells and, instead of a compensatory proliferation response, we detect even less EdU-positive cells with ageing, suggesting overall reduced proliferative capacity in the central retina (Fig 3C). These levels are further reduced after a 30-days chase (Fig 3D), suggesting that there are very few cells proliferating in the aged central retina. In the peripheral retina, where the proliferative CMZ resides, we see double the EdU-retaining cells in comparison to the central retina at 0-days chase (Fig 3C). Nevertheless, as in the central retina, proliferation in the peripheral retina (Fig 3 C and D) decrease with ageing, suggesting that there is no compensatory proliferation from the CMZ in response to the increased cell death with ageing. This also suggests that the CMZ has a finite proliferative capacity and the retina of the zebrafish does not continue to grow at the same rate throughout life. Finally, removing telomerase has no further effect on the already low and decreasing levels of proliferation with ageing in the retina. These results suggest that there is no compensatory proliferation in response to neurodegeneration in the aged zebrafish retina.

**Figure 3.**
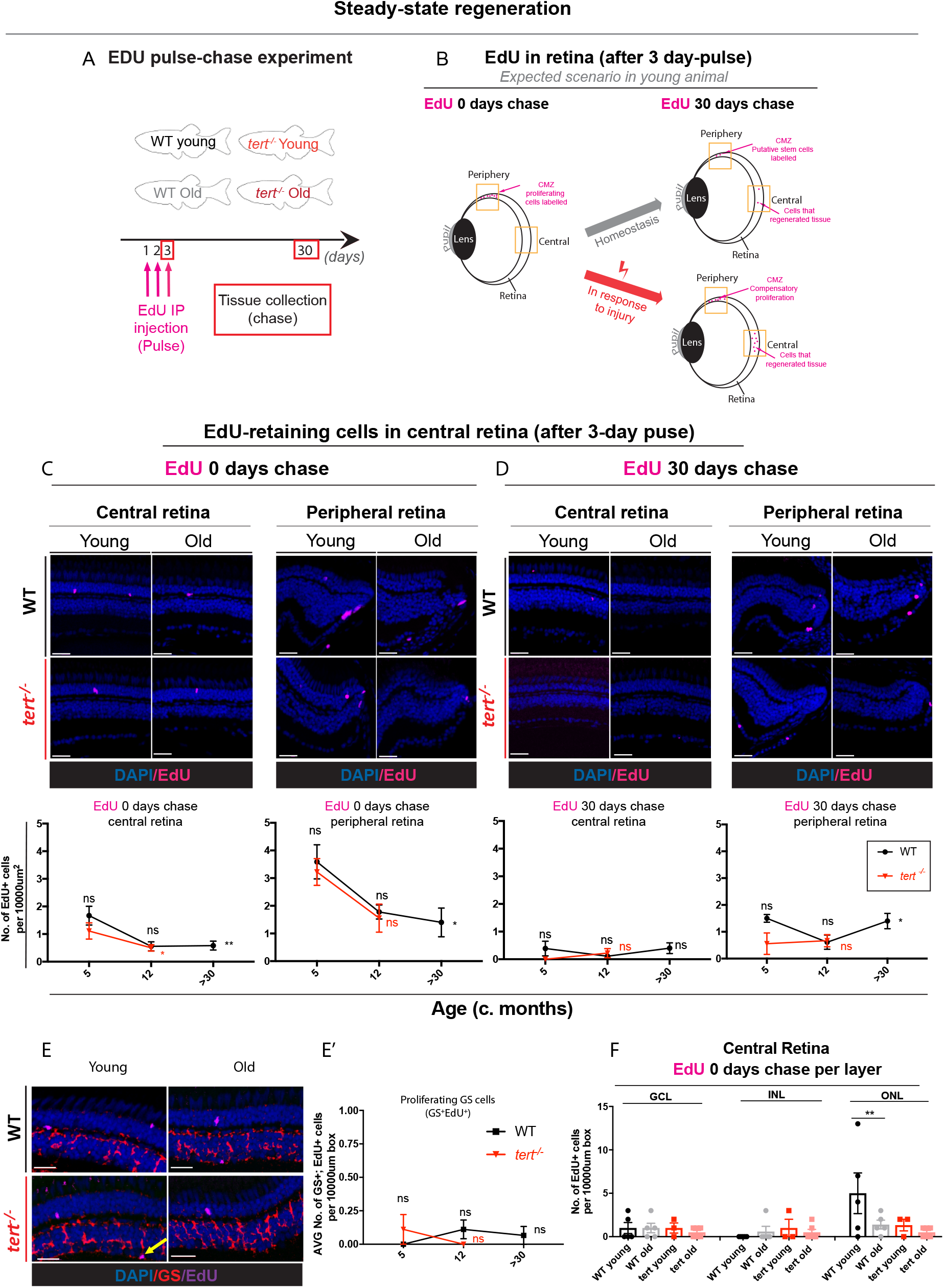
Aged zebrafish retina does not show signs of regeneration in response to spontaneous cell death and neuronal loss. (A) Schematic figure of the experimental design: 3-day pulse of EdU, by IP injection, followed by 0- or 30-day chase; (B) and respective hypothesised result. We anticipate that in a healthy young fish, the retina has some cells proliferating in the CMZ, which, over-time, will replace older cells in the central retina. In the case of injury, we anticipate that there will be elevated levels of proliferation in peripheral and central retina to replace the death cells. (C-D) The central and the peripheral retina immunolabeled with EdU (in purple), at (C) 0- and (D) 30-days chase, in both WT and *tert^-/-^*, at young (5 months) and old ages (>30 months in WT and 12 in *tert^-/-^*). Scale bars: 20μm. Graphs show quantifications of the number of EdU-retaining cells per area (10 000μm^2^), in the overall central and peripheral retina at (C) 0- and (D) 30-days chase. (E) The central retina immunolabeled with GS (Müller glia, in red) after a 3-day pulse of EdU, by IP injection, at 0days chase, in both WT and *tert^-/-^*, at young (5 months) and old ages (>30 months in WT and 12 in *tert^-/-^*). Scale bars: 20μm. (E’) Quantifications of the number of GS^+^; EdU^+^ cells per area (10 000μm^2^). Error bars represent SEM. N=3-6. CMZ: ciliary marginal zone. (F) Quantifications of the number of EdU-retaining cells per area (10 000μm^2^), per layer of the retina, at 0-days chase. Error bars represent SEM. N=3-6. P-value: * <0.05; ** <0.01; *** <0.001.

As proliferation of MG is the primary source of neurons after acute injury in fish ^12,36^, we sought to determine whether the EDU+ cells was indeed compensatory proliferation specifically within these cells. We co-labelled cells with the MG specific marker glutamine synthetase (GS), and counted the GS-positive; EdU-positive cells. We detect very few GS-positive; EdU-positive cells in the central retina in both WT and *tert^-/-^*, at any of the time-points characterised throughout their lifespan (Fig 3E, E’), suggesting MG are not proliferating in the central retina at old ages. We also do not observe PCNA expression in MG or signs of MG de-differentiation to progenitors^19^ in old age evidenced by a lack of Pax6 expression (Supplementary Fig 2). Thus, contrary to what has been reported to occur in response to acute damage^11–13^, our data show that age-related chronic cell death does not trigger proliferation of MG cells. Finally, rods can originate from rod-specific progenitors ^37,38^, which are found in the ONL and are derived from MG that slowly divide in the WT retina ^15,36,39,40^. To test whether there is any compensatory proliferation with ageing occurring from rod-specific progenitors found scattered throughout the ONL of the retina ^37,41^, we quantified levels of EdU positive cells in the ONL. Although we observe EDU+ cells in the central ONL of 5 months WT (likely to be rod precursors dividing), there are few detected in >30 months old WT. Once again, *tert^-/-^* mutants have no further effect (Fig 3F). Together, our data show that there is no compensatory proliferation in response to age-related degeneration in the zebrafish retina, by any of the known sources of regeneration and neurogenesis: CMZ, rod precursor cells or MG.

### Zebrafish vision declines with ageing, independently of telomerase

Loss of visual acuity and contrast sensitivity with advancing age in humans has been well documented ^6,7^ We report the presence of retinal neuron loss and tissue thinning in the aged retina, largely independently of telomerase. To determine whether the molecular and structural changes we observe have a pathological consequence on the retinal function, we tested whether zebrafish have impaired vision with ageing. Visual testing in the zebrafish has been used to screen for mutants with defects in retinal development and function ^42–45^, but are yet to be tested in the context of ageing. The optokinetic response (OKR) is a well-established assay to measure innate visual responses and provides readout of visual acuity ^42,46^ (Fig 4A). Our results show a decreased number of eye rotations per minute in aged fish (Fig 4B, B’ **and videos 1 and 2**), suggesting that zebrafish vision declines with ageing. As for most of the hallmark phenotypes of ageing described so far, (Supplementary Table 1), telomerase is not a limiting factor for visual acuity, since the *tert^-/-^* zebrafish do not display an accelerated reduced visual acuity at 5 months or 12 months of age, close to the end point of *tert^-/-^* life (Fig 4B, B’ **and videos 3 and 4**). As such, it does not appear that telomerase anticipates any of the hallmarks of zebrafish retina ageing tested here, unlike in other tissues^29^.

**Figure 4:**
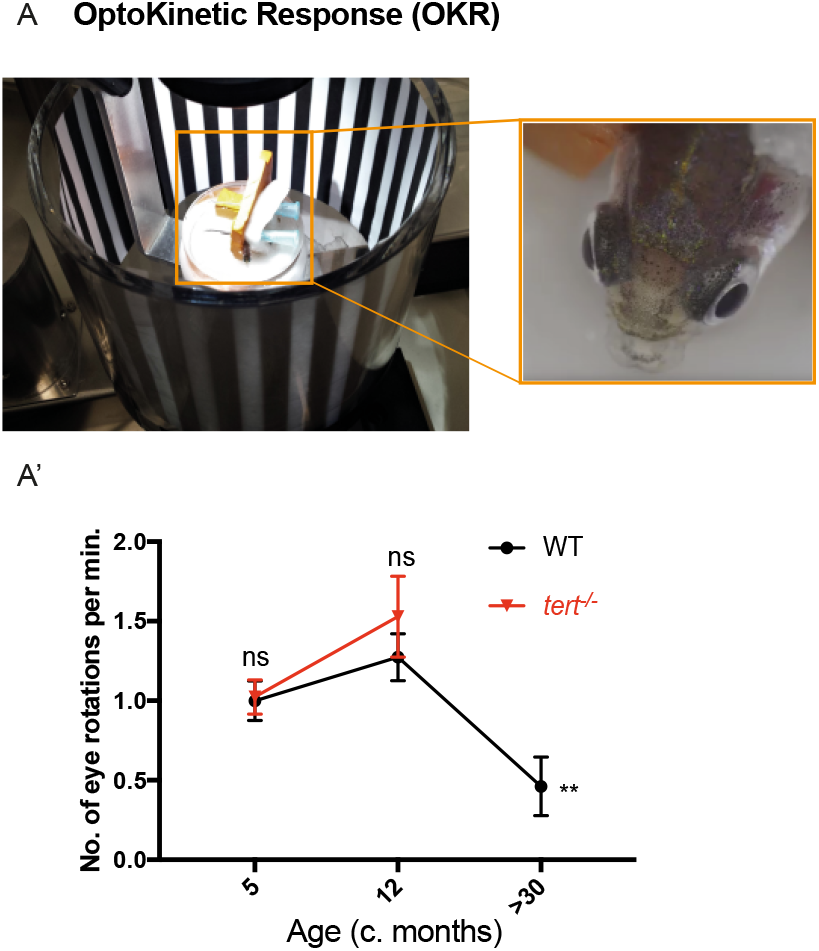
Zebrafish vision declines with ageing, independently of telomerase. (A) OKR assay was performed by immobilising the fish in between soft sponges, inside a petri dish containing water, placed in the centre of a rotation chamber. The walls of the rotation chamber had 0.8 mm-thick black and white stripes and the chamber was maintained at a constant velocity of 12 rpm throughout the experiment. (A’) The number of eye rotations per minute was manually quantified by video observation. Error bars represent the SEM. N=5-8. P-value: * <0.05; ** <0.01; *** <0.001.

### The zebrafish retina shows signs of gliosis with ageing

As we do not observe a MG regenerative response to neurodegeneration and vision loss, we wondered whether the retina was undergoing a gliotic response to damage. In the mammalian retina gliosis is a hallmark of retinal damage ^47^, but also commonly observed in the aged retina ^48^. MG respond to damage or injury in most, if not all, retinal degenerative diseases ^10^. The characteristic mammalian response is gliosis, whereby MG change shape and up-regulate structural proteins like glial fibrillary acidic protein (GFAP) to promote a neuroprotective effect ^9,10^. In the zebrafish retina, the initial regenerative response to acute damage is similar to mammalian gliosis ^18,49^, and gliosis can be aggravated if MG proliferation is blocked ^19^. Here, we identified MG using the well-described GS immunolabelling and characterised MG reactivity using the gliosis markers GFAP, Vimentin and Cralbp ^50,51^. We do not observe an overall loss of MG cell number in the central with ageing (Fig 5 A, A’), in contrast to the observed neuronal death observed in this region at the same stages (see Fig 2). Despite this, there is a significant change in the morphology of the MG cells in the WT aged retina. Aberrations in the aged MG cells include disruptions in the radial morphology along the synaptic IPL and basal lamina (Fig 5B-D), which are known hallmarks of gliosis in retina degeneration ^48^. Qualitative assessment further shows that while all young fish display long and aligned MG basal processes, 100% of the old fish show morphological disorganisation, a hallmark of gliosis. We also observed MG shape changes and changes in Vimentin and Cralbp expression at levels old ages in MG (Figure 5E-H). Thus, similarly to humans, chronic neurodegeneration with ageing elicits a MG gliotic response in the zebrafish retina rather than regeneration.

**Figure 5.**
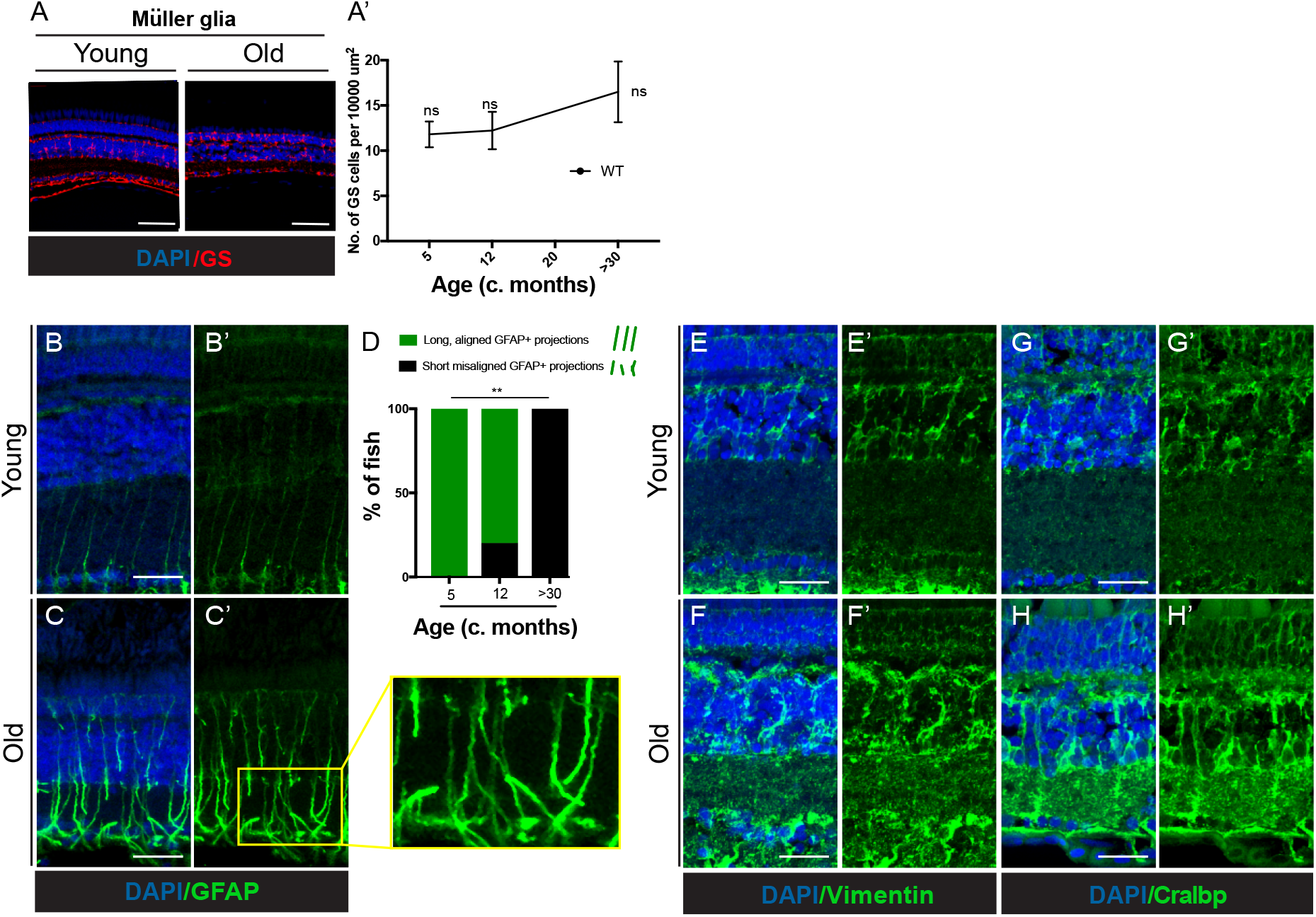
The zebrafish retina shows signs of gliosis with ageing. (A) The central retina immunolabeled with GS (Müller glia, in red) in WT fish, at young (5 months) and old ages (>30 months). Scale bars: 20μm. (A’) Quantification of the number of GS-positive cells (MG), per area (10 000 μm^2^). Error bars represent the SEM. N=3. (B,C) The central retina immunolabeled with GFAP (MG in green), in WT at young (5 months) and old ages (>30 months). (D) Quantifications of the percentage of fish presenting disorganised MG processes (gliosis-phenotype; inset) N = 6. P-value: * <0.05; ** <0.01; *** <0.001 (E,F) The central retina immunolabeled with Vimentin showing disorganised MG and upregulation of expression. (G,H) The central retina immunolabeled with Cralbp showing disorganised MG and upregulation of expression. Scale bars: 20μm.

Microglia, the innate immune cells found in the retina (reviewed in ^52^), are also key players in maintaining tissue homeostasis throughout life and part of the gliosis process. They are activated in many neurodegenerative diseases, with increased number at sites of damage, including in the photoreceptor layer in many forms of retinal degeneration (reviewed in ^48^). As we do not observe regeneration in the aged zebrafish retina in response to neuronal death, we asked whether there are alterations in the number of microglia (4C4-positive cells) found in the tissue. In contrast to what is observed in acute damage paradigms ^13,25^, we observe few microglia in the central retina, with no increase with ageing. Furthermore, microglia did not appear to be localised to regions of neuronal death, such as in photoreceptors or the INL (Supplementary Fig 3).

### Yap expression is maintained in MG throughout life

As we observe hallmarks of gliosis in MG cells and a lack of MG proliferation in response to age-associated chronic damage, we next asked whether gliotic MG have altered molecular profiles making them unable to proliferate in response to damage in the aged retina. Tgfβsignaling has been shown to differ between zebrafish regeneration and mouse gliosis response after acute damage, with Tgfβ3 being upregulated in regeneration and Tgfβ1 and 2 in gliosis^53^. However, we do not observe any difference in expression of any of the Tgfβisoforms in MG cells in the chronically aged zebrafish retina (Supplementary Fig 4). Hippo/Yap signaling has been identified as a signaling pathway that is altered after neuron degeneration and injury ^54–56^ within MG cells, and is required for MG regeneration^55^. Yap expression is also affected in reactive MG cells after damage^56^. We used antibodies against Yap, a downstream effector of Hippo signaling, to determine expression in MG throughout life. Yap is expressed in MG from 5dpf all the way through to end of life, including in gliotic MG cells (Fig 6). Yap is repressed from the nucleus after damage, suggesting that Yap activity may be required for the maintenance of MG proliferative capacity^56^. Consistent with this we do not observe any Yap positive nuclei at any stage suggesting that MG cells may retain the molecular composition to regenerate in response to retinal damage throughout lifespan.

**Figure 6.**
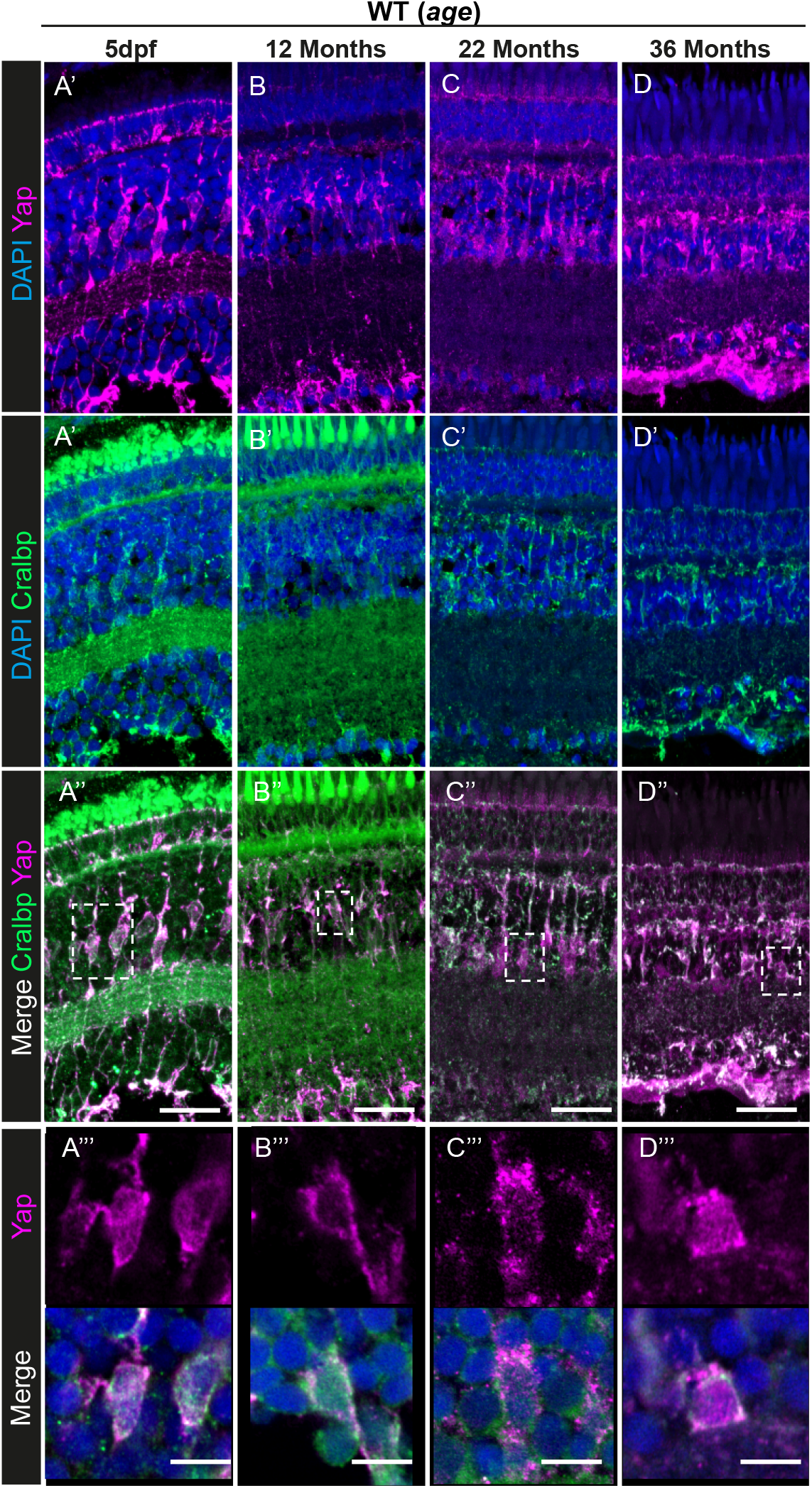
Yap expression is observed in the retinal MG throughout zebrafish lifespan. (A-D) The central retina immunolabeled with Yap (in magenta) and (A’-D’) Cralbp (MG cells, in green) in WT zebrafish at different ages (5 dpf, 12 months, 22 months, and 36 months). Scale bars: 20μm. (A’’’-D’’’) A single z-section from stacks showing a merge of DAPI, Yap and Cralbp staining with Yap expression specifically in MG cells at all stages. Scale bars: 3μm.

### Müller glia retain the ability to regenerate after acute damage until old age

Studies on retinal regeneration typically focus on the young adult retina and have not taken old age into account for the regenerative response. Several acute damage paradigms induce retinal neuron death and elicit a regenerative response in zebrafish ^36,49,57,58^. Chronic damage in the adult retina, specifically between 9-18 months of age, results in MG continuously reentering the cell cycle to proliferate and regenerate lost neurons, although they do show signs of chronic activation at later timepoints ^49^. However, it remains unclear whether MG retain their ability to proliferate and regenerate the retina throughout life, and whether gliotic MG, such as those often occurring in retinal disease or old age, also retain proliferative capacity to replace damaged neurons. As MG maintain Yap expression into old ages, we hypothesised that MG maintain the ability to regenerate in response to acute damage in the aged retina, despite not doing so after chronic degeneration. To test this, we used the light-damage model where aged *albino* zebrafish, at three stages of their lifecycle, were treated with light to elicit photoreceptor damage. To detect increased proliferation of MG in response to damage, a 3-day pulse of BrdU was performed, followed by a 28-days chase period (Fig 7A). After a 3-day pulse, BrdU should label both MG and their daughter cells, the newly formed progenitors (Fig 7B). However, since the BrdU staining dilutes in the rapidly dividing progenitor cells and MG only divide once, after a 28-days chase, the majority of the BrdU staining is retained in MG (Fig 7B). Our results show that light-lesion in aged retinas leads to a loss of photoreceptors, which is accompanied by a strong increase in microglia in the ONL, typical of photoreceptor degeneration in this damage model (Fig 7C, C’). Moreover, there are no differences in the incorporation of BrdU with ageing on any layer, at 72 hours post-light damage (hpL), when the initial regenerative response is mounted, or at 28 dpL, when the number of MG that initially re-entered the cycle can be identified (Fig 7D, D’). Both the unaltered immediate timing of MG response and the overall capacity to regenerate each neuronal layer are maintained with increased age. Finally, MG in the aged *albino* background also show a characteristic gliosis phenotype similar to WT (Supplementary Fig 5). Together these results suggest that the regenerative response remains intact throughout zebrafish lifespan and the molecular and morphological alterations in gliotic MG does not impact regenerative capacity in old age.

**Figure 7.**
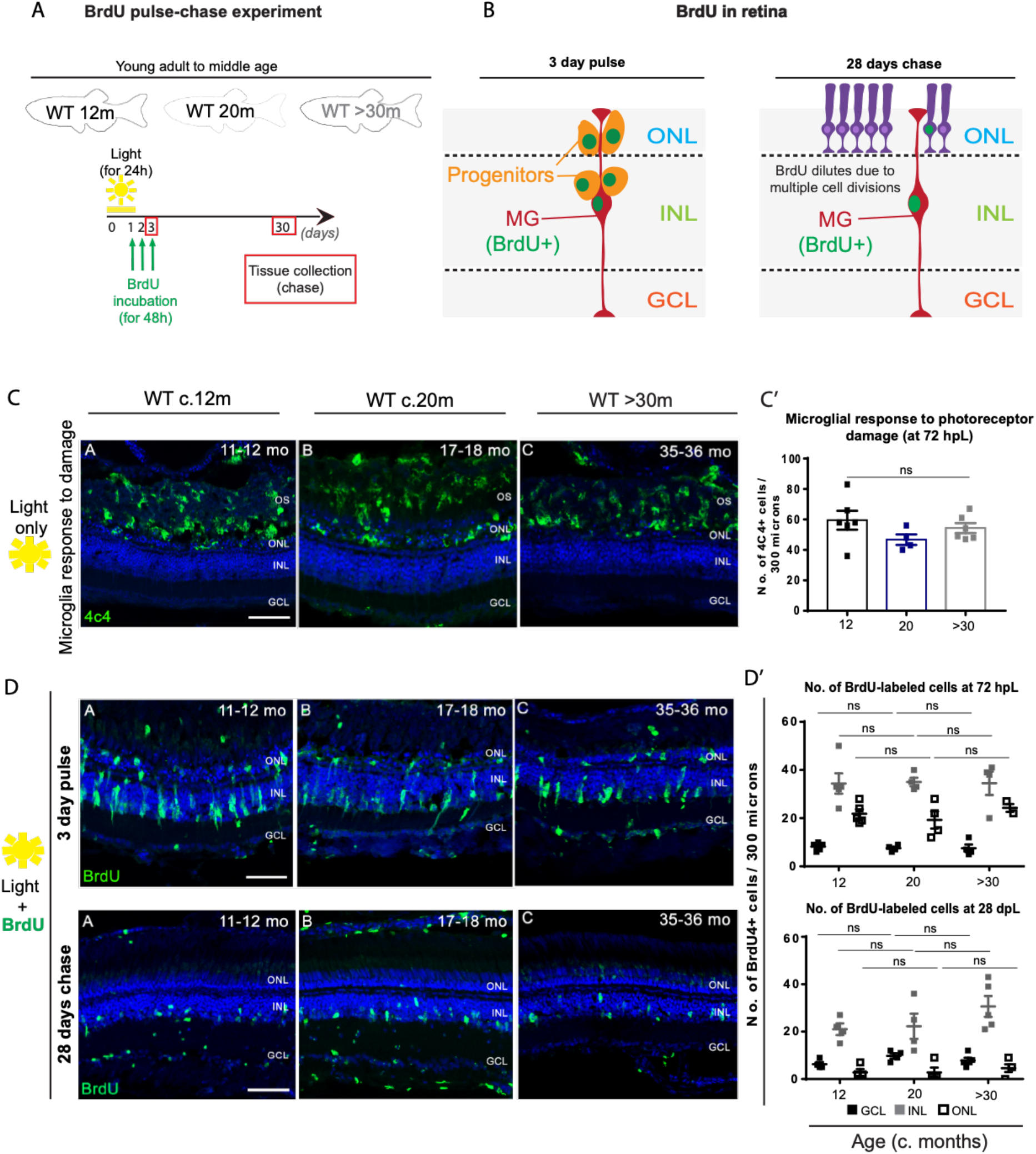
Zebrafish MG regenerative capacity upon acute damage is maintained until old age. . (A) Schematic image of the experimental design and (B) expected results. Fish were light-treated for 24 hours and BrdU incorporation occurred between 24-72 hrs (green), which allowed for BrdU to be washed out the proliferating progenitors, leaving only MG which reentered the cell cycle to be labelled. (C) The central retina labelled with 4C4 (microglia, in green), after light treatment, in young (c. 12 months), middle aged (c. 20 months) and old (>30 months) *albino* zebrafish. The majority of the microglia response to damage occurs within the outer segments (OS) and nuclear layer (ONL) of the retina, where a debris field is present due to photoreceptor degeneration. (C’) Quantification of the number of microglial cells which responded to photoreceptor damage in the different aged groups. (D, top) The central retina 72 hours after light treatment onset, immunolabeled with BrdU (proliferation, in green). (D’). Quantification of the number of cells proliferating in the ganglion cell layer (GCL, represented in black squares), inner nuclear layer (INL, represented in grey squares), and outer nuclear layer (ONL, represented in white squares) in each age group. (D, bottom) The central retina 28 days after light treatment onset. (D’). Quantification of the number of BrdU+ cells observed in the GCL, INL, and ONL of each aged group. Error bars represent the SEM. N=4-5. ns: nonsignificant. Scale bars: 20μm.

## DISCUSSION

Using a combination of cell labelling strategies throughout adulthood into old age, we show that the zebrafish retina retains its potential to regenerate in response to acute damage into old age. Therefore, the lack of compensatory proliferation in response to chronic, age-associated cell death in the ageing zebrafish retina is not due to a loss of capacity for MG to proliferate *perse*, but likely due to the absence or insufficient levels of the required stimuli for MG engagement. Indeed, contrasting to acute damage, we show that age-related retinal damage is insufficient to trigger compensatory proliferation by any of the known sources of regeneration and neurogenesis, namely the CMZ, rod precursor cells and MG. Moreover, we show that MG’s inability to heal the aged retina is not due to telomerase-dependent proliferative limits. Overall, instead of regeneration, zebrafish retina ageing leads to a gliotic response and loss of vision, reminiscent of human retinal ageing.

### Telomerase-dependent and -independent hallmarks of zebrafish retinal ageing

Previous work suggested that telomerase and telomere length are important for human retinal health. In particular, the retinal pigment epithelium (RPE) has been reported to have shorter telomeres than the neural retina, and it accumulates senescence over-time ^59^. Accordingly, reactivation of telomerase has been described to ameliorate symptoms of age-associated macular disease ^60,61^ and telomerase activators are currently in clinical trials (e.g. NCT02530255). However, retina homeostasis requires more than RPE maintenance. It requires a steady-state level of proliferation of different cell types involved in multiple aspects of retina function. As in other proliferative tissues ^29–31^, it would be expected that telomerase levels would influence zebrafish retina homeostasis. We therefore hypothesised that the zebrafish retina would degenerate in a telomerase-dependent manner, leading to vision loss.

However, our data suggest that most age-related changes in the zebrafish retina, including vision loss, are largely telomerase independent, since depletion of telomerase (*tert^-/-^*) does not accelerate or exacerbate any of these phenotypes. Telomere dysfunction is known to affect mostly highly proliferative tissues, reviewed in ^62^ and what our data show is that, in the region most affected by ageing phenotypes, the central retina, there is very little proliferation to start with. This suggests that replicative exhaustion of telomeres is not a limiting factor in the age-associated degeneration of the central retina. Instead, it is likely that chronic exposure to damaging agents such as oxidising UV radiation is the key driver of degeneration in the retina, as was proposed in the “Wear and Tear Theory” (reviewed in ^6^). However, we cannot exclude that telomere-associated damage may still be a contributing factor to the observed increased levels of DNA damage and cell death in the central retina. Telomeres are known to be damaged by oxidative stress and can act as sinks of DNA damage, irrespectively of length and levels of telomerase ^63,64^. In fact, this is a likely contributor to RPE damage with ageing ^65,66^, potentially explaining why depleting telomerase has no effect on the proliferation of the ONL, where photoreceptors reside, nor does it accelerate vision loss.

### Chronic vs. acute damage in MG responses to natural ageing

Our results show that zebrafish develop vision loss with ageing and that this is underpinned by retinal neurodegeneration and gliosis. This could seem counter-intuitive, given that it is well established that the zebrafish retina is capable of regenerating after acute damage. Therefore, the lack of regeneration after age-related cell death observed in this study suggest that there are critical differences between chronic ageing and repair after an acute injury. Further supporting this, we show that this lack of regeneration in the context of ageing is not due to an intrinsic inability of MG to proliferate *per se*, as we show they are still capable of doing so in response to an acute injury in old animals. Several tissues in the zebrafish retain the ability to regenerate throughout the lifespan of the zebrafish, including the fin and the heart ^67,68^. Likewise, the optic nerve crush paradigm has shown that there is successful recovery and function in the zebrafish retinotectal system ^69^, suggesting that regeneration is still possible at advanced age in the retina. The regenerative capacity of vertebrate tissues tends to decrease after repeated injury and as animals advance in age. This is likely due to reduced progenitor cell proliferation potential and subsequent differentiation ^70–73^. In the context of the central nervous system, this is aggravated by gliosis, a reactive change in glial cells in response to damage. In the mammalian retina, MG cells undergo gliosis in response to damage and in many retina degenerative diseases ^10^. However, in the zebrafish retina MG cells undergo the initial reactive gliotic response after acute damage but quickly shift to the regenerative pathway. Here we show that while we observe morphology alterations in MG cells in response to age-related neurodegeneration, MG in the aged retina retain their ability to regenerate in response to acute damage of photoreceptors, providing evidence that regenerative mechanisms in tissues that have been damaged or degeneration remains possible.

It is unclear if alternative damage paradigms will also lead to similar regenerative response and potential in the aged zebrafish retina. It also remains unclear what is the consequence of the gliotic response observed on retinal neurons in the aged retina, as gliosis is classically thought of as “Janus-faced”, with both pro- and anti-neuroprotective functions ^10^. Importantly, the kinetics of these MG morphology and molecular changes relative to neurodegeneration may provide further clues to the precise cellular breakdown in the retina causing widespread neuronal death and dysfunction. For instance, do the MG cells react first, thereby abandoning their neuronal support functions and precipitating neuronal damage, or do they purely respond to neuronal degeneration. Nevertheless, despite neuronal loss with ageing, MG cells maintain their numbers throughout ageing, suggesting that MG may be protected from age related death caused by the accumulation of DNA damage or free radicals. It remains unclear if there is a molecular mechanism inferring this protection or if MG gliosis is in itself protective to cell death. However, despite gliosis we show MG retain the ability to regenerate lost neurons after acute damage, which is critical for the potential to use regeneration as a therapy for damage in human retina in ageing or degenerative disease.

### Our proposed model: A molecular “tipping point” required to stimulate regeneration in ageing

Regeneration studies so far have relied on several damage paradigms, including phototoxic ^49^ and ouabain induced lesions ^74^, which induce rapid cell death post-insult. While acute damage models are suitable to explore the cellular and molecular mechanisms underpinning the regenerative potential of the retina, they do not allow testing whether this regenerative response is also occurring with natural ageing in the retina. “Natural ageing” and associated stress-induced neuronal death play out over months or even years and we now show that they do not elicit the same regenerative response. This may be because the slow degeneration is not producing a strong enough signal in a short amount of time to induce MG to undergo the cellular and molecular process of regeneration (Fig 8). It has been shown that the level of cell death can induce a differential response of MG cells. Whilst a large amount of rod death causes a regenerative response, small amounts do not ^26,75^. Furthermore, there may be distinct differences in signals released after apoptosis or necrosis in ageing vs damage models^76,77^. In support of this concept, recent work suggests that there may be key signalling differences underpinning the difference between a “regenerative” or a “reparative” response to injury ^53^. Thus, we propose that a molecular signal, or expression changes of such signal, will regulate the “tipping point” required to elicit a MG regenerative response in ageing. Research in the context of acute damage paradigms in the zebrafish retina has uncovered many of the molecular mechanisms regulating this regenerative response (reviewed in ^19, 37^). For instance, proliferation appears to be a key mechanism for the initiation of the regenerative response as blocking proliferation after damage in the zebrafish retina results in MG gliosis, and not regeneration, similar to humans ^16^. Moreover, age-associated degeneration of retinal neurons and their synapses, as we observed to occur in this study, may result in a loss of neurotransmitter release, such as GABA, which has been shown to facilitate the initiation of MG proliferation ^78^. Recently, it has been suggested that electrical stimulation may promote MG proliferation and expression of progenitor markers ^81^. Thus, dysregulation of neurotransmitters upon neurodegeneration could inhibit the key molecular pathways regulating regeneration. Alternatively, the initial inflammatory response has also been shown to be determinant for the repair process. Upon light-induced retinal damage, overexpression and subsequent release of TNF by apoptotic photoreceptors seems to induce MG proliferation ^18,79^. After acute damage, there is also an increased number of microglia in the retina ^74,80^ and they have been shown to be essential for retinal regeneration ^81,82^. In contrast, microglia numbers do not increase with ageing and therefore may compromise the regenerative response. Nonetheless, it is important to consider that the available immunohistochemistry techniques to identify microglia number has a few limitations. Namely, there may be a spike in microglia number in the aged retina that is rapidly resolved and missed at our timepoints or microglia may undergo apoptosis similar to retinal neurons. Thus, we cannot exclude the possibility that microglial cells are involved in the observed retinal degenerations or play a critical role in the lack of a regenerative response observed in the aged retina. Identifying this signal will be critical to stimulate regeneration in humans as using acute light damage is unlikely to be therapeutically feasible. Finally, our data suggest that, as YAP expression is maintained throughout lifespan, manipulating the Hippo/Yap pathways in the aged retina may be a suitable model to treat retina degenerations into old age.

**Figure 8:**
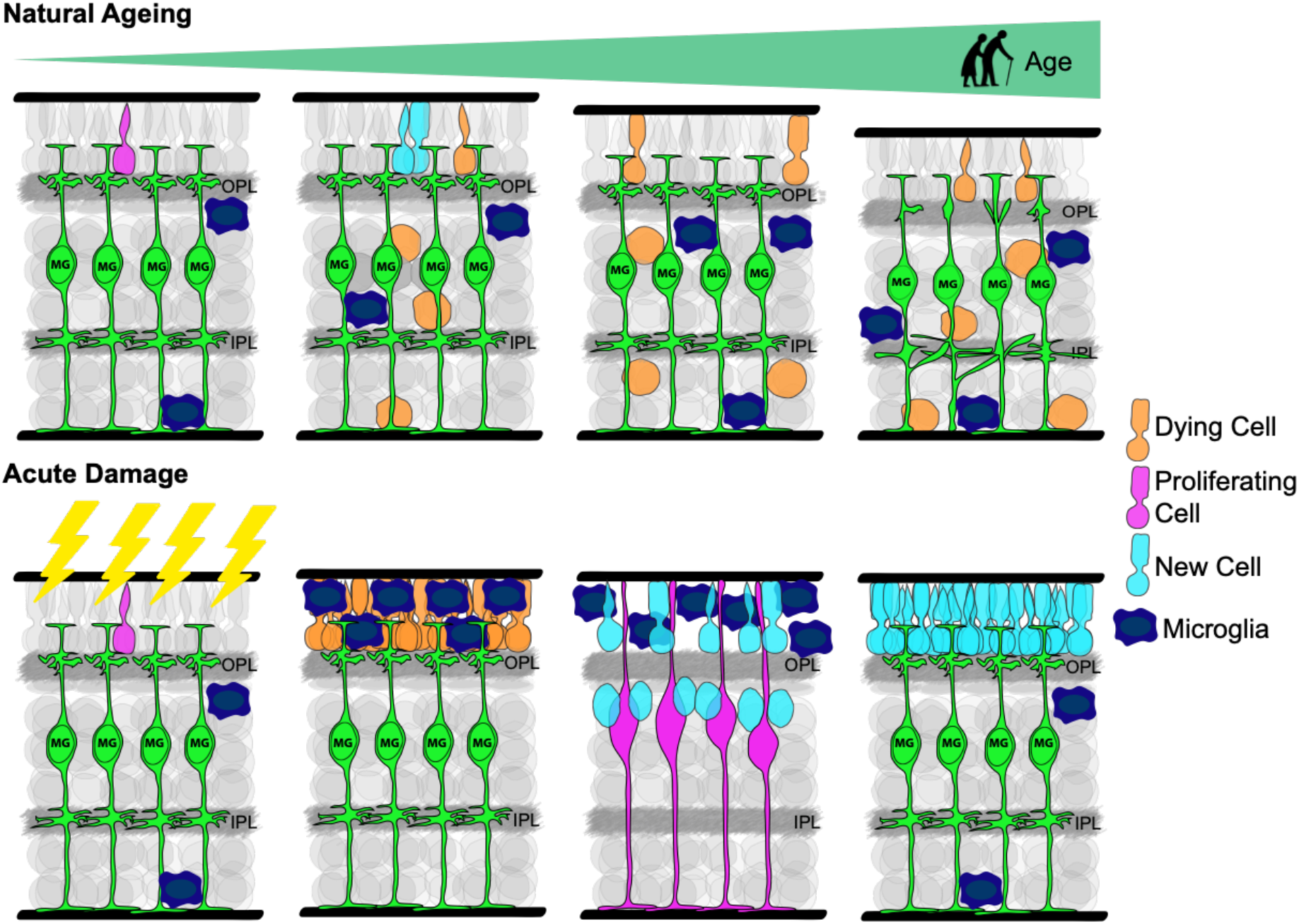
Our proposed model: A molecular “tipping point” required to stimulate regeneration in ageing. Schematic figure with the working model. Top: In natural ageing, zebrafish retina undergoes degeneration characterised by increased cell death and neuronal loss (represented in orange). Our current findings show that the proliferation levels are low in young ages and decrease even more with advancing age (represented in magenta). Moreover, the number of microglia, a key play in retina regeneration, is maintained stable throughout lifespan (represented in dark blue). The lack of MG proliferation in response to chronic, age-related damage seems to lead to retina thinning with ageing, where death cells are not replaced. In fact, instead of proliferating in response age-related chronic damage, zebrafish MG cells undergo a gliotic-like phenotype, similarly to humans. Despite maintaining Yap expression until old ages, MG cells do not stimulate regeneration of the aged zebrafish retina. Bottom: In contrast with age-associated chronic damage, in response to acute damage where there is a great number of dying cells, MG proliferates, generating new neuronal cells to replace the dying ones (represented in light blue). Here, we show that zebrafish gliotic MG cells retain their regenerative capacity in response to acute light damage until old ages (>30 months). These findings highlight key differences in the MG response to chronic versus acute damage and show that gliotic MG cells can be stimulated to repair damaged neurons in the old zebrafish retina.

## Conclusions

Our work demonstrates that, in the context of age-induced neuronal degeneration, the MG in the zebrafish retina react by undergoing a response more akin to gliosis, rather than regeneration. This resembles what occurs in the aged human retina as well as in many human retinal degenerative diseases. Importantly, we identify key differences between chronic versus acute damage and show that gliotic MG cells can be stimulated to repair damaged neurons in the old zebrafish retina.

## MATERIALS AND METHODS

### Zebrafish husbandry

Zebrafish were maintained at 27-28°C, in a 14:10 hour (h) light-dark cycle and fed twice a day. The OKR was performed in the UCL Institute of Ophthalmology and the phototoxic lesions and regeneration experiments were performed at Wayne State University School of Medicine (USA). All other experiments were performed in the University of Sheffield. All animal work was approved by local animal review boards, including the Local Ethical Review Committee at the University of Sheffield (performed according to the protocols of Project Licence 70/8681) and the Institutional Animal Care and Use Committee at Wayne State University School of Medicine (performed according to the protocols of IACUC-19-02-0970).

### Zebrafish strains, ages and sex

Three strains of adult zebrafish (*Danio rerio*) were used for these studies: wild-type (WT; AB strain), *tert^-/-^* (*tert^AB/hu3430^*) and *albino (slc45a2^b4/b4^). tert^-/-^* zebrafish is a premature model of ageing and therefore, age and die earlier than the naturally aged zebrafish. While *tert^-/-^* fish have a lifespan of 12-20 months, WT fish typically die between 36-42 months of age ^29,30^. In order to study age-related phenotypes in the zebrafish retina, here we used young (5 months) WT and *tert^-/-^* fish, alongside with middle aged (12 months) WT fish, and old WT and *tert^-/-^* fish. ‘Old’ was defined as the age at which the majority of the fish present age-associated phenotypes, such as cachexia, loss of body mass and curvature of the spine. These phenotypes develop close to the time of death and are observed at >30 months of age in WT and at >12 months in *tert^-/-^*^29,30^. In addition, we used adult *albino* zebrafish for retinal regeneration studies (described in detail below). Importantly, none of the animals included in this study displayed visible morphological alterations in the eyes (e.g. cataracts). Whenever possible, males where chosen to perform the experiments.

### OKR assay

Fish were anaesthetised in 4% tricaine methanesulfonate (MS-222; Sigma-Aldrich) and placed in a small bed-like structure made of sponge, located inside a small petri dish containing fish water. During the experiment, fish were maintained still by the strategic use of needles that sustained the sponges close to the fish so that the fish could not move. The petri dish was then placed inside a rotation chamber with black and white striped walls (8mm-thick stripes). After fish recovered from anaesthesia, the trial began, and the walls of the chamber started rotating at 12rpm (for 1min to the left side followed by 1min to the right side). Eye movements were recorded using a digital camera throughout the experiment. After the experiment, the number of eye rotations per minute was measured by video observation and manually counting. The counting was performed blindly by two independent researchers. In the end, the results were normalised for the WT young from the same day / batch, in order to control for different days of experiments.

### Intense light-damage paradigm with BrdU incorporation

A photolytic damage model in adult *albino* zebrafish was utilised to destroy rod and cone photoreceptors and elicit a regenerative response ^26^. Briefly, adult *albino* zebrafish were dark-adapted for 10 days prior to a 30 min exposure to ~100,000 lux from a broadband light source. Next, fish were exposed to ~10,000 lux of light from four, 250 W halogen lamps for 24 hrs. Following 24 hrs of light treatment, fish were transferred to a 1L solution containing 0.66 g of NaCl, 0.1 g Neutral Regulator (SeaChem Laboratories, Inc. Stone Mountain, GA, USA), and 1.5 g BrdU (5mM; B5002; Sigma-Aldrich) for 48 hrs. This timeframe for BrdU incubation was based on previous studies in order to label MG cell-cycle re-entry ^49^. Following a 48 hour incubation in BrdU, the fish were split into two groups: one group was euthanised by an overdose of 2-Phenoxyenthanol and eyes were processed for immunohistochemistry as described below; the second group returned to normal husbandry conditions for an additional 25 days (or 28 days after light onset) prior to euthanasia and tissue collection. During this time, BrdU incorporation dilutes in actively dividing cells, allowing for clear visualization of the number of MG in the INL that only divide a single time. It also serves as an indirect measure of regenerative capacity (i.e. an equal loss of BrdU-positive cells at 28 dpL between experimental groups would indicate similar numbers of progenitor cell divisions earlier in the regenerative process).

### Tissue preparation: paraffin-embedded sections and cryosections

Adult fish were culled by overdose of MS-222, followed by confirmation of death. Whole fish or dissected eyeballs were then processed for paraffin-embedded sections or for cryosections, as follows:

Paraffin-embedded sections. Whole fish were fixed in in 4% paraformaldehyde (PFA) buffered at pH 7.0, at 4°C for 48-72h, decalcified in 0.5M EDTA at pH 8.0 for 48-72h, and embedded in paraffin by the following series of washes: formalin I (Merck & Co, Kenilworth, NJ, USA) for 10min, formalin II for 50min, ethanol 50% for 1h, ethanol 70% for 1h, ethanol 95% for 1h 30min, ethanol 100% for 2h, ethanol 100% for 2h 30min, 50:50 of ethanol 100%: xilol for 1h 30min, xylene I for 3h, xylene II for 3h, paraffin I for 3h and paraffin II for 4h 30min. Paraffin-embedded whole fish were then sliced in sagittal 4μm-thick or coronal 16μm-thick sections, using a Leica TP 1020 cryostat.

Cryopreservation and cryosections. Dissected eyeballs were fixed in 4% PFA at 4°C, overnight (ON). Then, they were washed in cold 1x PBS and immersed in 30% sucrose in phosphate-buffered saline (PBS), ON at 4°C, for cryopreservation. Single cryopreserved eyeballs were then embedded in mounting media - optimal cutting temperature compound (OCT, VWR International), snap-frozen at −80°C, and stored at −20°C until cryosectioning. Cryosections were sliced at a 13μm thickness using a Leica Jung Frigocut cryostat or a Leica CM1860 cryostat.

### Immunohistochemistry (IHC)

Before immunofluorescence staining, cryosections were hydrated in PBS at room temperature (RT) for 10min, and paraffin-embedded sections were deparaffinised and hydrated as follows: histoclear (Scientific Laboratory Supplies, Wilford, Nottingham, UK) 2x for 5min, followed by ethanol 100% 2x for 5min, ethanol 90% for 5min, ethanol 70% for 5min, and distilled water 2x for 5min. After antigen retrieval in 0.01M citrate buffer at pH 6.0 for 10min, the sections were permeabilised in PBS 0.5% Triton X-100 for 10min and blocked in 3% bovine serum albumin (BSA), 5% Goat Serum (or Donkey Serum), 0.3% Tween-20 in PBS, for 1h. The slides were then incubated with the primary antibody at 4°C ON. After washes in PBS 0.1 % Tween-20 (3x 10min) to remove excess to primary antibody, the sections were incubated with secondary antibody at RT for 1h. Finally, the slides were incubated in 1μg/ml of 4’,6-diamidino-2-phenylindole (DAPI, Thermo Fisher Scientific) at RT for 10min, washed in PBS 1x, and mounted with vectashield (Vector Laboratories, Burlingame, CA, USA). The primary and secondary antibodies used in this study are described in Table 1 and Table 2, respectively.

**Table 2.**
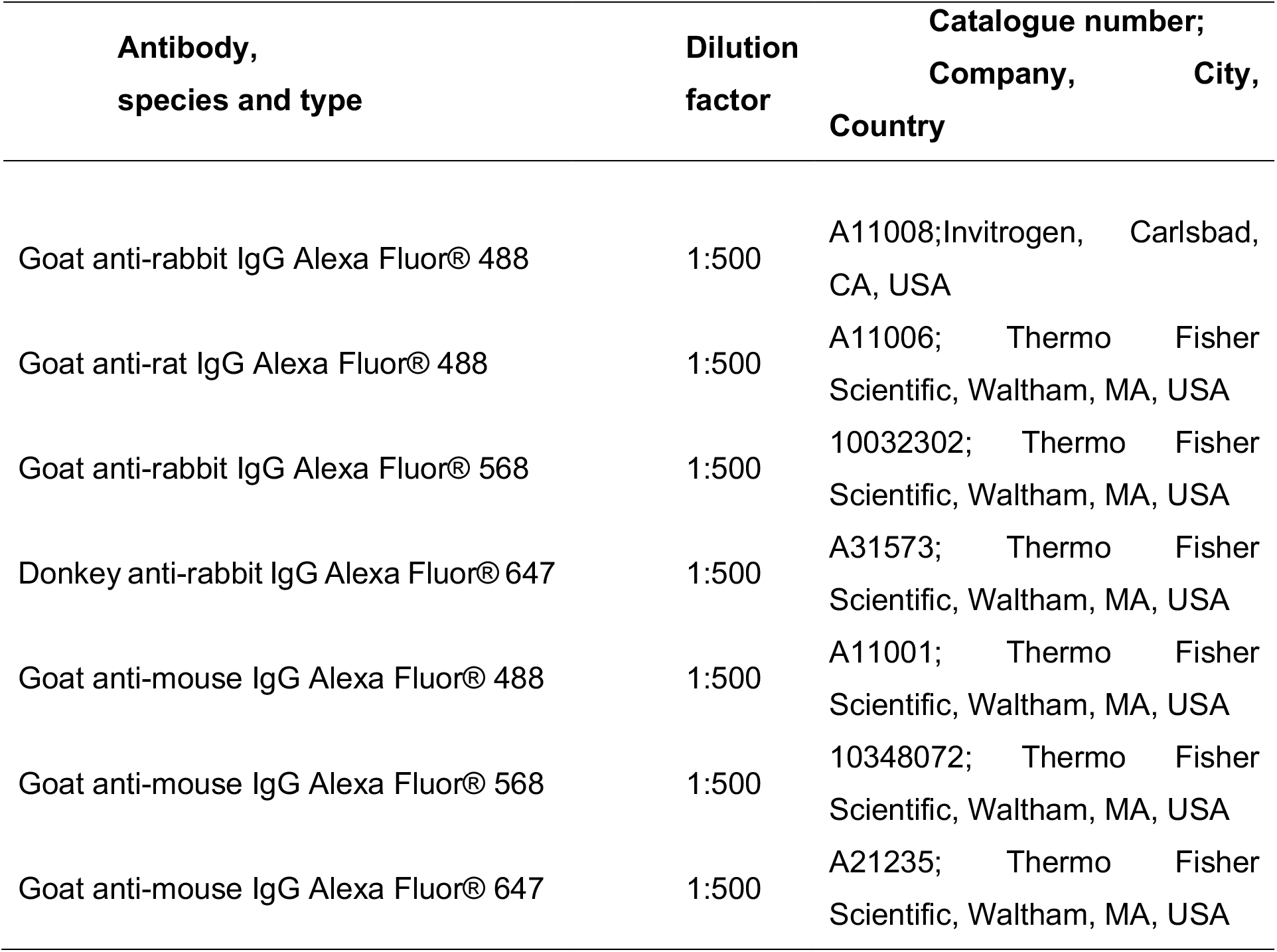
Secondary antibodies used for immunostaining.

### Terminal deoxynucleotidyl transferase dUTP nick end labelling (TUNEL) staining

In paraffin-embedded sections, TUNEL was performed using the *In Situ Cell Death Detection Kit, Fluorescein* (Merck & Co), following the manufacturer’s instructions. Briefly, after deparaffinisation, hydration, antigen retrieval and permeabilization, as described above, the slides were incubated in enzyme and label solution (1:10) at 37°C for 1h. The slides were then washed in 1x PBS (2x 10min) before blocking and incubation with primary antibody.

### 5-Ethynyl-2′-deoxyuridine (EDU) labelling

EdU labelling was detected using the Click-iT® EdU Imaging Kit (Thermo Fisher Scientific), following manufacturer’s instructions. Briefly, fish were injected with 5μl of 10mM EdU diluted in dimethyl sulfoxide (DSMO), by intraperitoneal (IP) injection, for 3 consecutive days (3-day pulse). In order to differentiate proliferating cells and low-proliferative EdU-retaining cells, the fish were separated into two groups: 0-day chase and 30-days chase groups. The fish from the first group were culled 2h30min after the last injection of Edu, whereas the fish from the second group were culled 30 days after the last injection of EdU. After culling, whole fish were processed for paraffin-embedded sections as described above. In order to detect EdU labelling in paraffin-embedded sections, the slides were deparaffinised, hydrated, underwent antigen retrieved, were permeabilised and washed in 1x PBS. The slides were incubated in freshly made EdU-labelling solution (per 1ml of solution: 860 1x Click-iT®EdU reaction buffer, 40 CuSO4, 2.5 Alexa Fluor®647 azide working solution and 100 10x EdU reaction buffer additive) at RT for 30min. Finally, the slides were washed in 1x PBS before blocking and incubation with primary antibody (ON, at 4°C). The incubation in secondary antibody was performed as previously described.

### Imaging and quantifications

Paraffin-embedded sections were imaged by epifluorescence microscopy, using a DeltaVision microscope with a 40x oil objective. Cryosections were imaged by laser scanning confocal imaging, using a Leica SP5 microscope, Nikon A1 Confocal microscope or a Zeiss 900 Airyscan 2 confocal, with a 40x oil objective. In either paraffin or cryosections, multiple 0.2- 0.6μm thick z-stacks were acquired in order to capture the whole retina region. For each staining, a total of 4 images were taken per retina, 2 from central and 2 from peripheral retinal. The central retina was defined to be the centre point between opposing CMZs (A minimum of ~1000μm from the periphery), while images of the peripheral retina contain the limits of the retina, including the CMZ. The peripheral retina was defined as the tissue directly adjacent to the CMZ.

In order to quantify the alteration in the staining patterns, a z-projection was generated and three boxes of 100×100μm were drawn in each field of view. The total number of positive cells was then manually counted for each labelling. Rhodopsin, ZO1, ribeye and GFAP staining were an exception to this, for which a qualitative assessment was performed instead. To do so, structural and morphological defects were identified as follows. For the rhodopsin staining, as young WT retinas usually display long and aligned outer segments, all short and/or misaligned outer segments were considered defective. For the ZO1 staining, the average number of breaks in the ZO1-labelled membrane per animal was quantified. The average of breaks in the WT young animals was used as a reference, and any fish presenting an average number of breaks bellow this average was considered to present defects in the membrane. For the ribeye staining, young retinas usually present two distinguished layers of pre-synaptic ribbons. Thus, staining where the two layers of pre-synaptic ribbons are not distinguished, was considered defective. GFAP staining usually reveals long and aligned MG processes in young WT retinas, and therefore short and/or misaligned MG processes are considered gliotic. Through this qualitative assessment it was calculated the percentage of fish per group presenting structural and morphological defects. Finally, raw images were used for quantification purposes. The images were then processed with Adobe Illustrator 21.0.2 for display purposes.

### Statistical analysis

Statistics were performed using the GraphPad Prism v7.00. Normality was assessed by Shapiro-Wilk test. For normally distributed data unpaired t-test was used to compare 2 data points and one-way ANOVA followed by Bonferroni post-hoc test was used to compare more than 2 data points. For non-normally distributed data, Mann-Whitney test and Kruskal-Wallis test followed by Dunn’s post-hoc test were used instead. Two-way ANOVA was used in order to compare more than 2 data points in 2 groups different groups (genotypes). Chi-square was performed to analyse structure and morphological changes in the retina based on qualitative assessment, having into account the number of animals per group displaying defects versus not displaying defects. A critical value for significance of p<0.05 was used throughout the analysis.

## ACKNOWLEDGEMENTS

The authors would like to thank Alex McGowen for 4c4 and Teresa Nicholson for RibeyeA for their generous gifts of antibodies. Part of this work was supported by grants from the National Institutes of Health (NEI R01EY026551 and R21EY031526 to RT). Histology and imaging core resources were supported by vision core grants (P30EY04068) and an unrestricted grant from Research to Prevent Blindness to the Department of Ophthalmology, Visual and Anatomical Sciences (RT). This work was also generously funded by a University of Sheffield PhD studentship to RRM, a Sheffield University Vice Chancellor’s Research Fellowship and a Wellcome Trust/Royal Society Sir Henry Dale Fellowship (UNS35121) to CMH and a Welcome Trust Seed Award (210152/Z/18/Z) and a BBSRC David Phillips Fellowship (BB/S010386/1) to RBM.

## Supplementary Data

**Supplementary Table 1:**
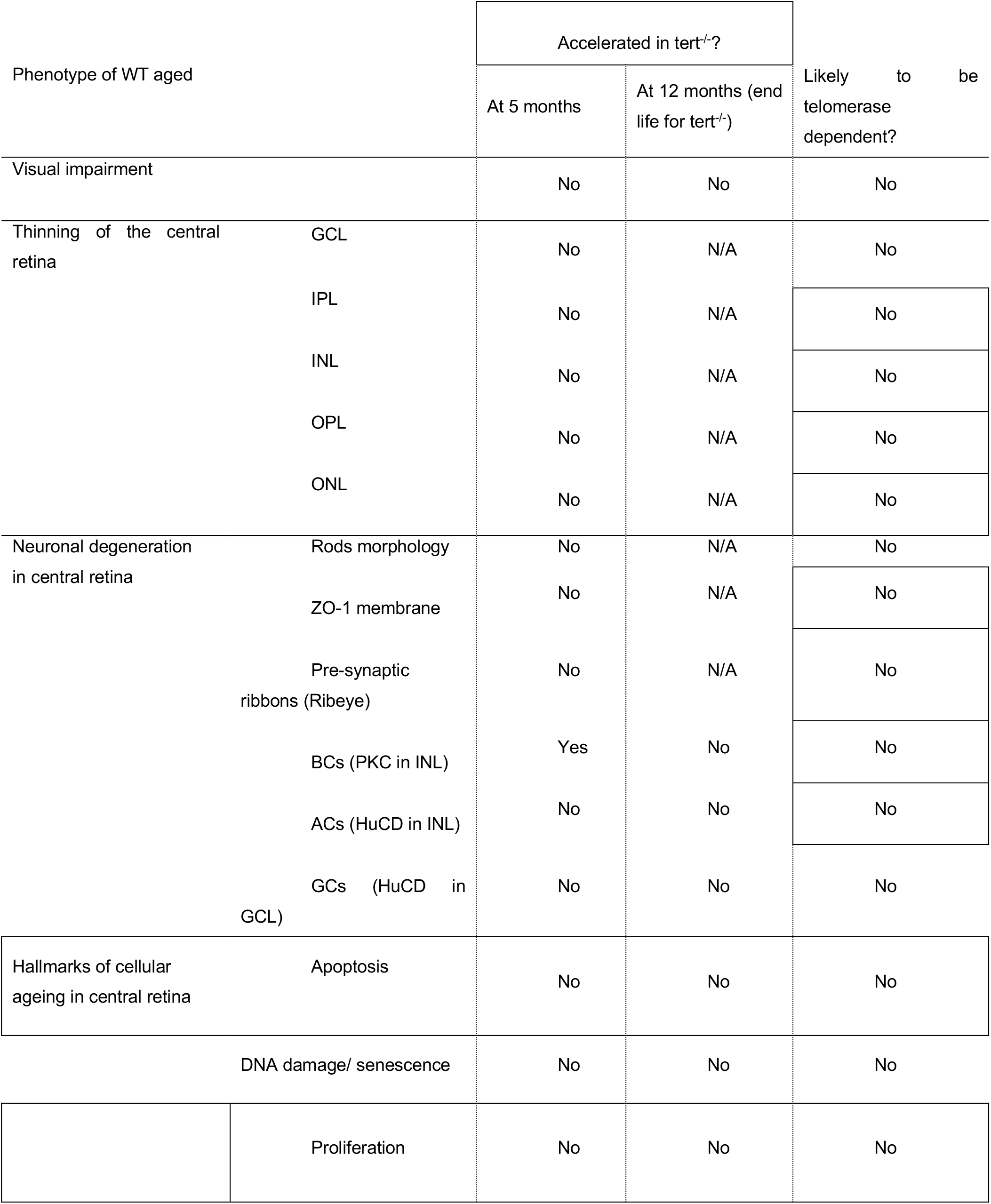
Summary of the phenotypes observed in the aged zebrafish retina, and which phenotypes are telomerase-dependent or independent.

**Supplementary Fig 1:**
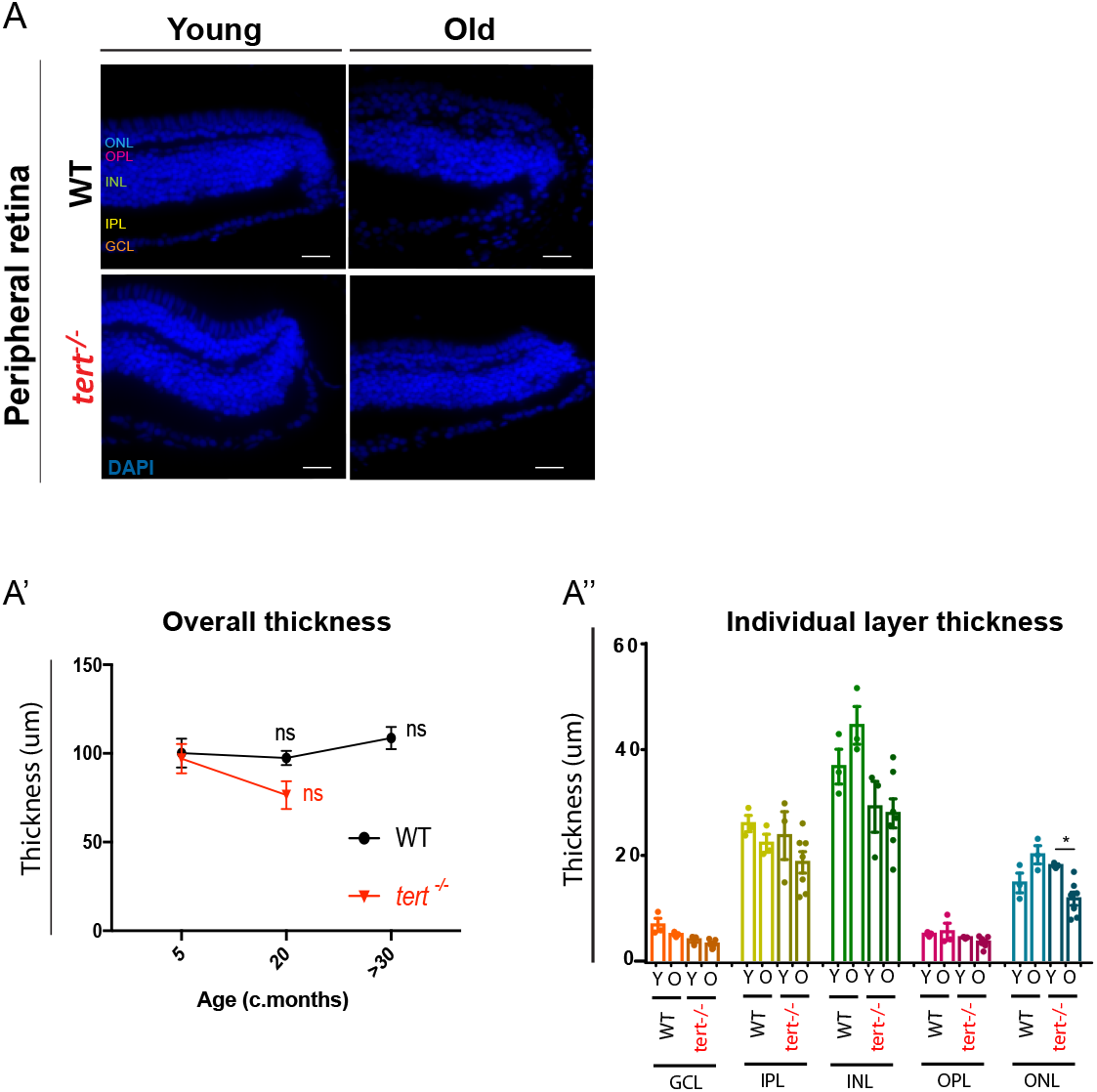
Key hallmarks of ageing in the zebrafish peripheral retina. (A) Peripheral retina thickness in both WT and *tert^-/-^*, at young (5 months) and old ages (>30 months in WT and 20 months in *tert^-/-^*). Scale bars: 20μm. (A’-A’’) Quantifications of the peripheral retina thickness cells per area (10 000μm^2^), (A’) in the overall retina and (A’’) per layer of the retina. GCL: ganglion cell layer; IPL: inner plexiform layer; INL: inner nuclear layer; OPL: outer plexiform layer; ONL: outer nuclear layer. Error bars represent the SEM. N=3. P-value: * <0.05; ** <0.01; *** <0.001.

**Supplementary Figure 2:**
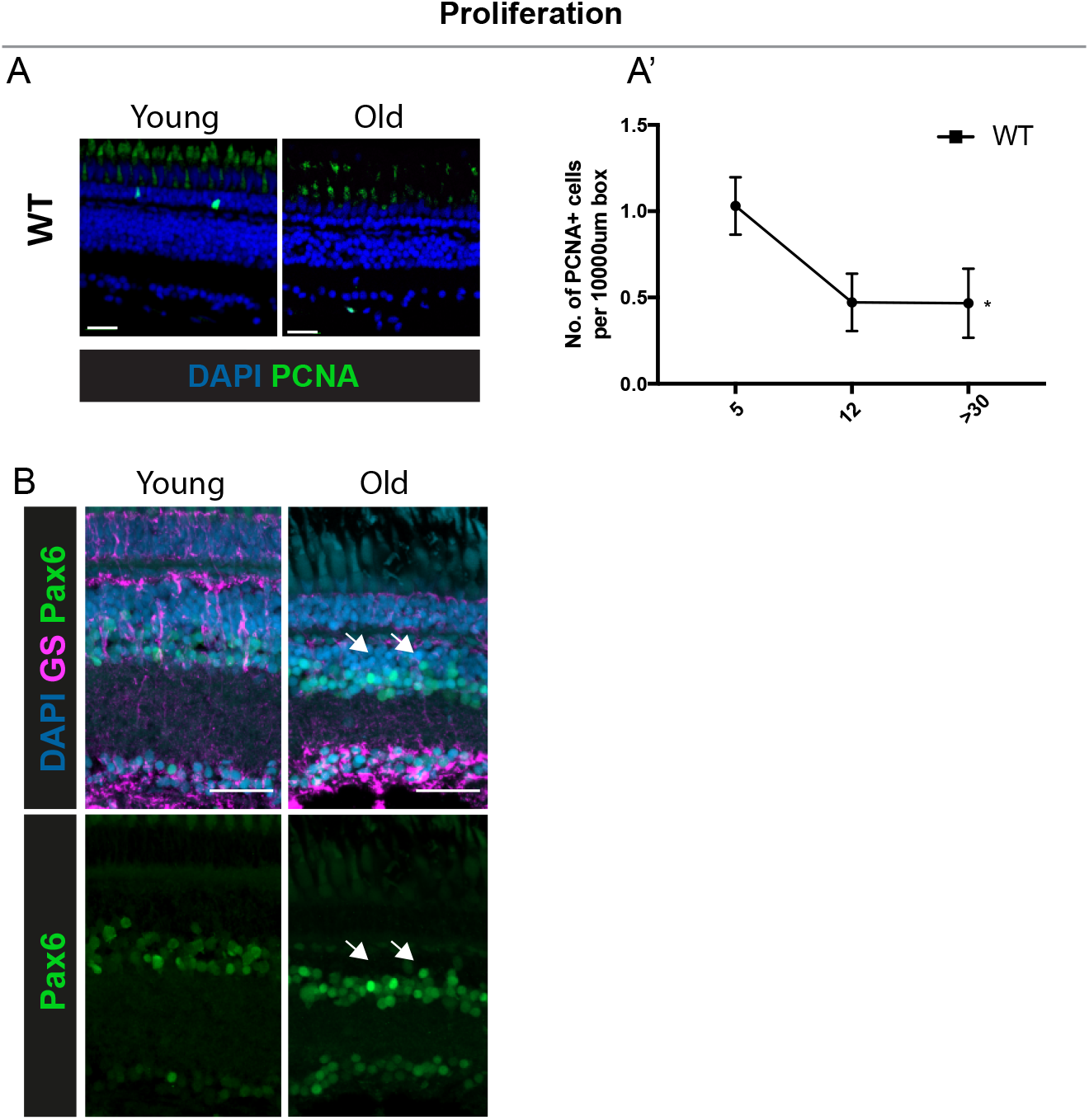
Lack of MG proliferation and dedifferentiation in the aged zebrafish retina. (A) The central retina immunolabeled with PCNA (cells proliferating, in green) in WT at young (5 months) and old ages (>30 months). Scale bars: 20μm. (A’) Quantifications of the number of PCNA^+^ cells per area (10 000μm^2^). Error bars represent the SEM. N=6-12. P-value: * <0.05; ** <0.01; *** <0.001. (B) We do not observe Pax6+ MG cells or progenitors in the aged zebrafish retina (arrows). The central retina co-immunolabeled with GS (MG cells, in magenta) and Pax6 (in green). N= 5.

**Supplementary Figure 3:**
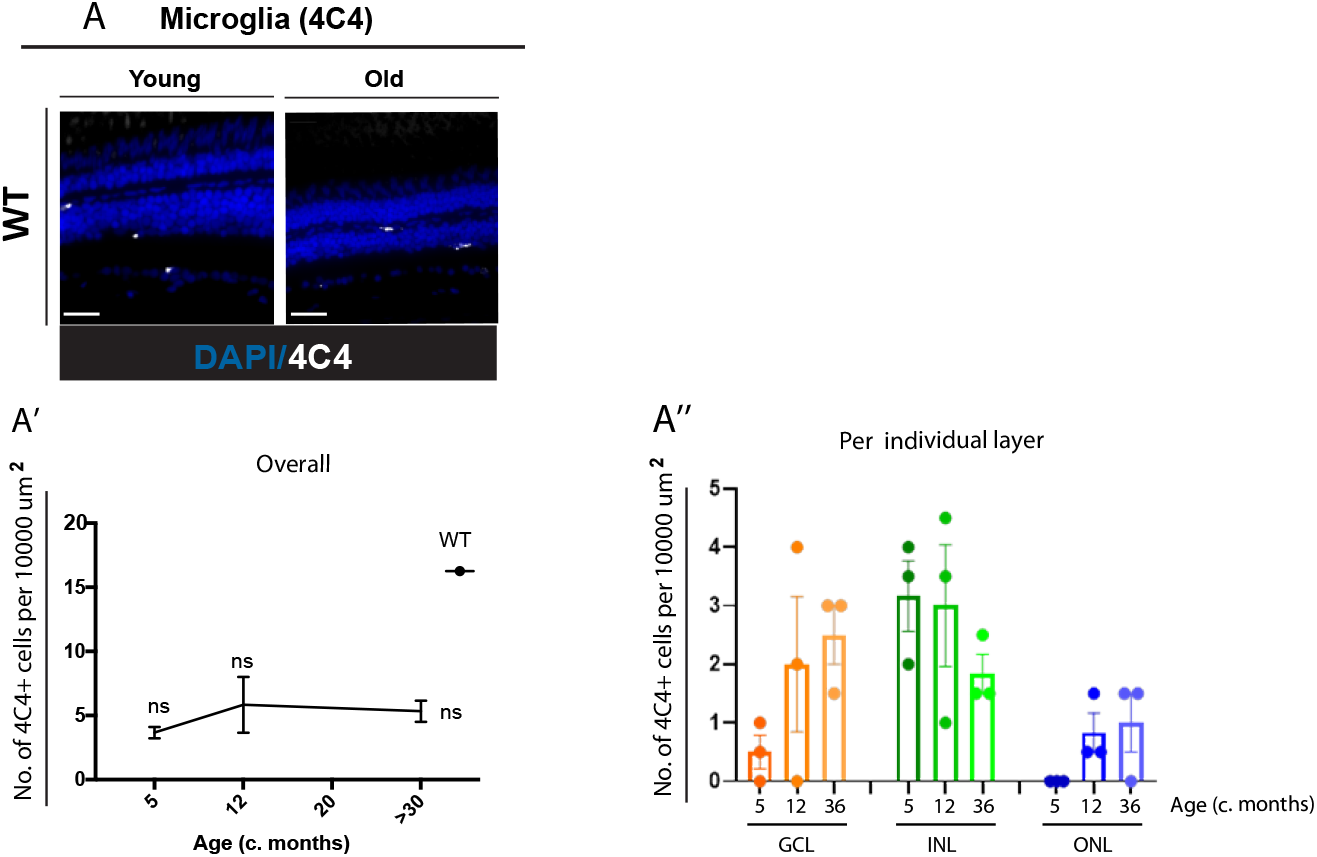
Microglia do not increase in number or distribution in the ageing zebrafish retina. (A) The central retina immunolabeled with 4C4 (microglia, in white) in WT zebrafish, from young (5 months) to old ages (>30 months). Scale bars: 20μm. (A’, A’’) Quantification of the number of 4C4-positive cells (microglia) per area (10 000 μm2) in the (A’) overall retina and (A’’) per layer of the retina. GCL: ganglion cell layer; INL: Inner nuclear layer; ONL: outer nuclear layer. Error bars represent the SEM. N=3. P-value: *<0.05; ** <0.01; *** <0.001.

**Supplementary Figure 4:**
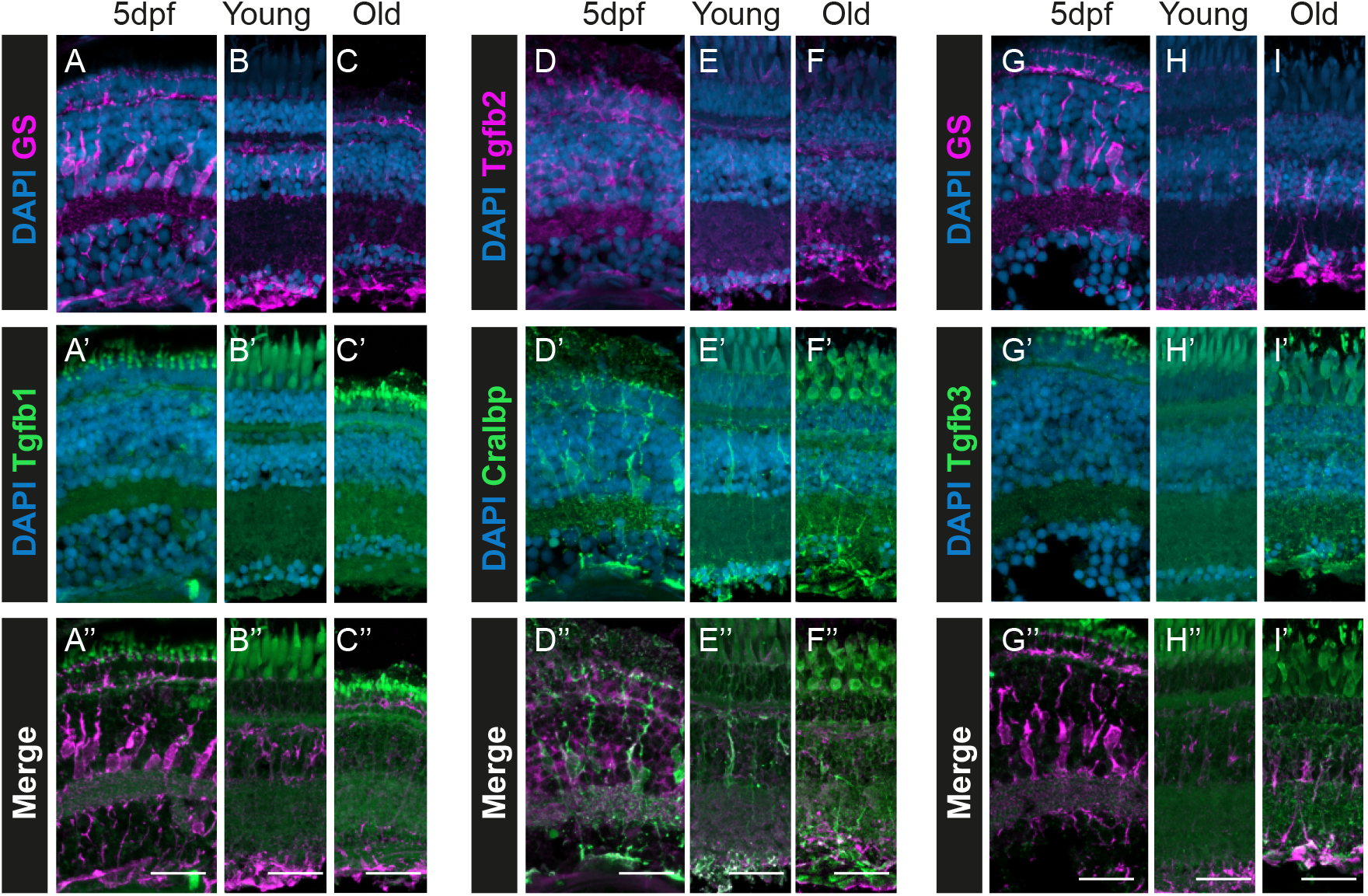
Expression of Tgfβ1, 2 and 3 in the zebrafish retina with ageing. We do not observe Tgfβ expression in MG cells in the old retina. The central retina co-immunolabeled with (A-C) GS (MG cells, in magenta) and (A’-C’) Tgfβ1 (in green); (D-F) Tgfβ2 (in magenta) and (D’-F’) Cralbp (MG cells, in green); and GS (MG cells, in magenta) and Tgfβ3 (in green), in WT zebrafish, from young to old ages. Scale bars: 20μm.

**Supplementary Figure 5:**
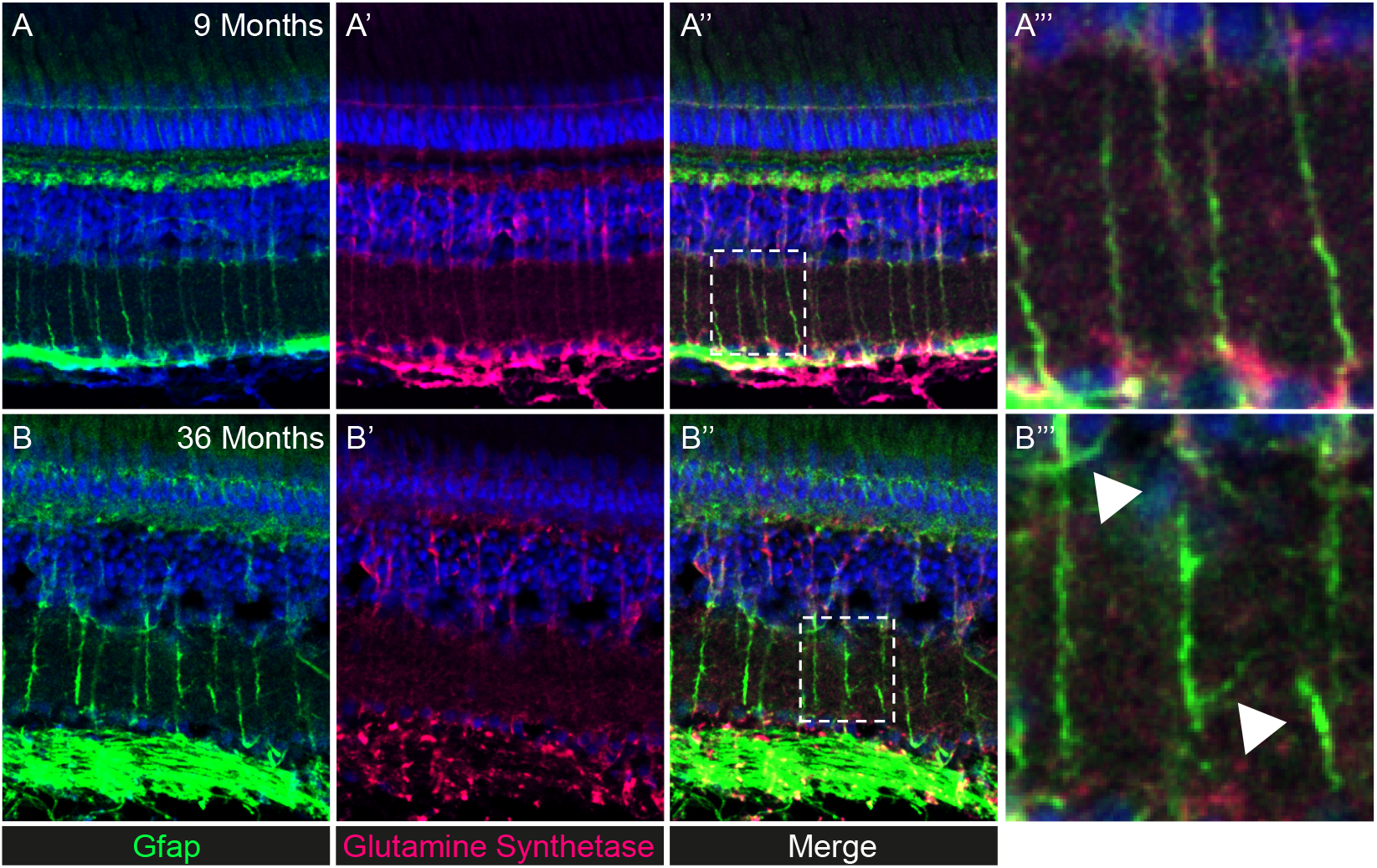
Mutant *albino* zebrafish display gliosis with ageing in MG similar to AB wildtype zebrafish. The central retina co-immunolabeled with (A, B) GFAP (reactive MG, in green) and (A’, B’) glutamine synthetase (MG, in magenta) in *albino* zebrafish at (A) 9 and (B) 36 months of age. (A’’’, B’’’) Insets of (A’’, B’’) merge staining showing disorganisation of MG processes in the IPL.

## REFERENCES

1. Chakravarthy, U. et al. Clinical risk factors for age-related macular degeneration: a systematic review and meta-analysis. BMC Ophthalmol 10, 31 (2010).

2. FJ, S. AGE NORMS OF REFRACTION AND VISION. Arch Ophthalmol 43, 466–481 (1950).

3. Weale, R.A. Senile changes in visual acuity. Trans Ophthalmol Soc U K 95, 36–8 (1975).

4. Arden, G.B. & Jacobson, J.J. A simple grating test for contrast sensitivity: preliminary results indicate value in screening for glaucoma. Invest Ophthalmol Vis Sci 17, 23–32 (1978).

5. Skalka, H.W. Effect of age on Arden grating acuity. Br J Ophthalmol 64, 21–3 (1980).

6. Marshall, J. The ageing retina: physiology or pathology. Eye (Lond) 1 (Pt 2), 282–95 (1987).

7. Salvi, S.M., Akhtar, S. & Currie, Z. Ageing changes in the eye. Postgrad Med J 82, 581–7 (2006).

8. Al-Ubaidi, M.R., Naash, M.I. & Conley, S.M. A perspective on the role of the extracellular matrix in progressive retinal degenerative disorders. Invest Ophthalmol Vis Sci 54, 8119–24 (2013).

9. Reichenbach, A. & Bringmann, A. New functions of Muller cells. Glia 61, 651–78 (2013).

10. Bringmann, A. et al. Cellular signaling and factors involved in Muller cell gliosis: neuroprotective and detrimental effects. Prog Retin Eye Res 28, 423–51 (2009).

11. Goldman, D. Muller glial cell reprogramming and retina regeneration. Nat Rev Neurosci 15, 431–42 (2014).

12. Fausett, B.V. & Goldman, D. A role for alpha1 tubulin-expressing Muller glia in regeneration of the injured zebrafish retina. J Neurosci 26, 6303–13 (2006).

13. Fischer, A.J., McGuire, C.R., Dierks, B.D. & Reh, T.A. Insulin and fibroblast growth factor 2 activate a neurogenic program in Muller glia of the chicken retina. J Neurosci 22, 9387–98 (2002).

14. Lenkowski, J.R. & Raymond, P.A. Muller glia: Stem cells for generation and regeneration of retinal neurons in teleost fish. Prog Retin Eye Res 40, 94–123 (2014).

15. Raymond, P.a., Barthel, L.K., Bernardos, R.L. & Perkowski, J.J. Molecular characterization of retinal stem cells and their niches in adult zebrafish. BMC developmental biology 6, 36 (2006).

16. Thummel, R., Kassen, S.C., Montgomery, J.E., Enright, J.M. & Hyde, D.R. Inhibition of Muller glial cell division blocks regeneration of the light-damaged zebrafish retina. Dev Neurobiol 68, 392–408 (2008).

17. Wan, J. & Goldman, D. Retina regeneration in zebrafish. Curr Opin Genet Dev 40, 41–47 (2016).

18. Thomas, J.L., Ranski, A.H., Morgan, G.W. & Thummel, R. Reactive gliosis in the adult zebrafish retina. Exp Eye Res 143, 98–109 (2016).

19. Thummel, R. et al. Characterization of Muller glia and neuronal progenitors during adult zebrafish retinal regeneration. Exp Eye Res 87, 433–44 (2008).

20. Eastlake, K. et al. Comparison of proteomic profiles in the zebrafish retina during experimental degeneration and regeneration. Sci Rep 7, 44601 (2017).

21. Jadhav, A.P., Roesch, K. & Cepko, C.L. Development and neurogenic potential of Muller glial cells in the vertebrate retina. Prog Retin Eye Res 28, 249–62 (2009).

22. Ooto, S. et al. Potential for neural regeneration after neurotoxic injury in the adult mammalian retina. Proc Natl Acad Sci U S A 101, 13654–9 (2004).

23. Jorstad, N.L. et al. Stimulation of functional neuronal regeneration from Muller glia in adult mice. Nature 548, 103–107 (2017).

24. Karl, M.O. et al. Stimulation of neural regeneration in the mouse retina. Proc Natl Acad Sci U S A 105, 19508–13 (2008).

25. Jorstad, N.L. et al. STAT Signaling Modifies Ascl1 Chromatin Binding and Limits Neural Regeneration from Muller Glia in Adult Mouse Retina. Cell Rep 30, 2195–2208 e5 (2020).

26. Thomas, J.L., Nelson, C.M., Luo, X., Hyde, D.R. & Thummel, R. Characterization of multiple light damage paradigms reveals regional differences in photoreceptor loss. Exp Eye Res 97, 105–16 (2012).

27. Van Houcke, J. et al. Extensive growth is followed by neurodegenerative pathology in the continuously expanding adult zebrafish retina. Biogerontology 20, 109–125 (2019).

28. Fu, J., Nagashima, M., Guo, C., Raymond, P.A. & Wei, X. Novel Animal Model of Crumbs-Dependent Progressive Retinal Degeneration That Targets Specific Cone Subtypes. Invest Ophthalmol Vis Sci 59, 505–518 (2018).

29. Carneiro, M.C. et al. Short Telomeres in Key Tissues Initiate Local and Systemic Aging in Zebrafish. PLoS genetics 12, e1005798 (2016).

30. Henriques, C.M., Carneiro, M.C., Tenente, I.M., Jacinto, A. & Ferreira, M.G. Telomerase is required for zebrafish lifespan. PLoS Genet 9, e1003214 (2013).

31. Anchelin, M. et al. Premature aging in telomerase-deficient zebrafish. Dis Model Mech 6, 1101–12 (2013).

32. Bednarek, D. et al. Telomerase Is Essential for Zebrafish Heart Regeneration. Cell Rep 12, 1691–703 (2015).

33. Easter, S.S., Jr. & Nicola, G.N. The development of vision in the zebrafish (Danio rerio). Dev Biol 180, 646–63 (1996).

34. Eriksson, U. & Alm, A. Macular thickness decreases with age in normal eyes: a study on the macular thickness map protocol in the Stratus OCT. Br J Ophthalmol 93, 1448–52 (2009).

35. Lo, S.N.a.A.C.Y. Protecting the Aging Retina. in Neuroprotection (ed. Ho, R.C.- C.C.a.Y.-S.) (2019).

36. Bernardos, R.L., Barthel, L.K., Meyers, J.R. & Raymond, P.A. Late-stage neuronal progenitors in the retina are radial Muller glia that function as retinal stem cells. J Neurosci 27, 7028–40 (2007).

37. Johns, P.R. & Fernald, R.D. Genesis of rods in teleost fish retina. Nature 293, 141–2 (1981).

38. Julian, D., Ennis, K. & Korenbrot, J.I. Birth and fate of proliferative cells in the inner nuclear layer of the mature fish retina. J Comp Neurol 394, 271–82 (1998).

39. Otteson, D.C., D’Costa, A.R. & Hitchcock, P.F. Putative stem cells and the lineage of rod photoreceptors in the mature retina of the goldfish. Dev Biol 232, 62–76 (2001).

40. Nelson, S.M., Frey, R.A., Wardwell, S.L. & Stenkamp, D.L. The developmental sequence of gene expression within the rod photoreceptor lineage in embryonic zebrafish. Dev Dyn 237, 2903–17 (2008).

41. Stenkamp, D.L. Development of the Vertebrate Eye and Retina. Prog Mol Biol Transl Sci 134, 397–414 (2015).

42. Brockerhoff, S.E. et al. A behavioral screen for isolating zebrafish mutants with visual system defects. Proc Natl Acad Sci U S A 92, 10545–9 (1995).

43. Neuhauss, S.C. et al. Genetic disorders of vision revealed by a behavioral screen of 400 essential loci in zebrafish. J Neurosci 19, 8603–15 (1999).

44. Gross, N.E., Aizman, A., Brucker, A., Klancnik, J.M., Jr. & Yannuzzi, L.A. Nature and risk of neovascularization in the fellow eye of patients with unilateral retinal angiomatous proliferation. Retina 25, 713–8 (2005).

45. Lessieur, E.M. et al. Ciliary genes arl13b, ahi1 and cc2d2a differentially modify expression of visual acuity phenotypes but do not enhance retinal degeneration due to mutation of cep290 in zebrafish. PLoS One 14, e0213960 (2019).

46. Tappeiner, C. et al. Visual acuity and contrast sensitivity of adult zebrafish. Front Zool 9, 10 (2012).

47. Bringmann, A. & Wiedemann, P. Muller glial cells in retinal disease. Ophthalmologica 227, 1–19 (2012).

48. Telegina, D.V., Kozhevnikova, O.S. & Kolosova, N.G. Changes in Retinal Glial Cells with Age and during Development of Age-Related Macular Degeneration. Biochemistry (Mosc) 83, 1009–1017 (2018).

49. Ranski, A.H., Kramer, A.C., Morgan, G.W., Perez, J.L. & Thummel, R. Characterization of retinal regeneration in adult zebrafish following multiple rounds of phototoxic lesion. PeerJ 6, e5646 (2018).

50. Garcia, M. & Vecino, E. Role of Muller glia in neuroprotection and regeneration in the retina. Histol Histopathol 18, 1205–18 (2003).

51. Verardo, M.R. et al. Abnormal reactivity of muller cells after retinal detachment in mice deficient in GFAP and vimentin. Invest Ophthalmol Vis Sci 49, 3659–65 (2008).

52. Cragnolini, A.B. et al. Regional brain susceptibility to neurodegeneration: what is the role of glial cells? Neural Regen Res 15, 838–842 (2020).

53. Conedera, F.M. et al. Diverse Signaling by TGFbeta Isoforms in Response to Focal Injury is Associated with Either Retinal Regeneration or Reactive Gliosis. Cell Mol Neurobiol (2020).

54. Hamon, A. et al. Retinal Degeneration Triggers the Activation of YAP/TEAD in Reactive Muller Cells. Invest Ophthalmol Vis Sci 58, 1941–1953 (2017).

55. Hamon, A. et al. Linking YAP to Muller Glia Quiescence Exit in the Degenerative Retina. Cell Rep 27, 1712–1725 e6 (2019).

56. Rueda, E.M. et al. The Hippo Pathway Blocks Mammalian Retinal Muller Glial Cell Reprogramming. Cell Rep 27, 1637–1649 e6 (2019).

57. Cameron, D.A. Cellular proliferation and neurogenesis in the injured retina of adult zebrafish. Vis Neurosci 17, 789–97 (2000).

58. Hanovice, N.J. et al. Regeneration of the zebrafish retinal pigment epithelium after widespread genetic ablation. PLoS Genet 15, e1007939 (2019).

59. Drigeard Desgarnier, M.C. et al. Telomere Length Measurement in Different Ocular Structures: A Potential Implication in Corneal Endothelium Pathogenesis. Invest Ophthalmol Vis Sci 57, 5547–5555 (2016).

60. Dow, C.T. & Harley, C.B. Evaluation of an oral telomerase activator for early age-related macular degeneration - a pilot study. Clin Ophthalmol 10, 243–9 (2016).

61. Rowe-Rendleman, C. & Glickman, R.D. Possible therapy for age-related macular degeneration using human telomerase. Brain research bulletin 62, 549–53 (2004).

62. Henriques, C.M. & Ferreira, M.G. Consequences of telomere shortening during lifespan. Curr Opin Cell Biol 24, 804–8 (2012).

63. Fumagalli, M. et al. Telomeric DNA damage is irreparable and causes persistent DNA-damage-response activation. Nat Cell Biol 14, 355–65 (2012).

64. Hewitt, G. et al. Telomeres are favoured targets of a persistent DNA damage response in ageing and stress-induced senescence. Nat Commun 3, 708 (2012).

65. Honda, S., Hjelmeland, L.M. & Handa, J.T. Oxidative stress--induced single-strand breaks in chromosomal telomeres of human retinal pigment epithelial cells in vitro. Invest Ophthalmol Vis Sci 42, 2139–44 (2001).

66. Wang, J. et al. Photosensitization of A2E triggers telomere dysfunction and accelerates retinal pigment epithelium senescence. Cell Death Dis 9, 178 (2018).

67. Itou, J., Kawakami, H., Burgoyne, T. & Kawakami, Y. Life-long preservation of the regenerative capacity in the fin and heart in zebrafish. Biol Open 1, 739–46 (2012).

68. Pinto-Teixeira, F. et al. Inexhaustible hair-cell regeneration in young and aged zebrafish. Biol Open 4, 903–9 (2015).

69. Van Houcke, J. et al. Successful optic nerve regeneration in the senescent zebrafish despite age-related decline of cell intrinsic and extrinsic response processes. Neurobiol Aging 60, 1–10 (2017).

70. Collins, C.A., Zammit, P.S., Ruiz, A.P., Morgan, J.E. & Partridge, T.A. A population of myogenic stem cells that survives skeletal muscle aging. Stem Cells 25, 885–94 (2007).

71. Janzen, V. et al. Stem-cell ageing modified by the cyclin-dependent kinase inhibitor p16INK4a. Nature 443, 421–6 (2006).

72. Nave, K.A. Neuroscience: An ageing view of myelin repair. Nature 455, 478–9 (2008).

73. Kirschner, D.A. et al. Rapid assessment of internodal myelin integrity in central nervous system tissue. J Neurosci Res 88, 712–21 (2010).

74. Mitchell, D.M., Sun, C., Hunter, S.S., New, D.D. & Stenkamp, D.L. Regeneration associated transcriptional signature of retinal microglia and macrophages. Sci Rep 9, 4768 (2019).

75. Montgomery, J.E., Parsons, M.J. & Hyde, D.R. A novel model of retinal ablation demonstrates that the extent of rod cell death regulates the origin of the regenerated zebrafish rod photoreceptors. J Comp Neurol 518, 800–14 (2010).

76. Bergmann, A. & Steller, H. Apoptosis, stem cells, and tissue regeneration. Sci Signal 3, re8 (2010).

77. Nelson, C.M. et al. Tumor necrosis factor-alpha is produced by dying retinal neurons and is required for Muller glia proliferation during zebrafish retinal regeneration. J Neurosci 33, 6524–39 (2013).

78. Rao, M.B., Didiano, D. & Patton, J.G. Neurotransmitter-Regulated Regeneration in the Zebrafish Retina. Stem Cell Reports 8, 831–842 (2017).

79. Iribarne, M., Hyde, D.R. & Masai, I. TNFalpha Induces Muller Glia to Transition From Non-proliferative Gliosis to a Regenerative Response in Mutant Zebrafish Presenting Chronic Photoreceptor Degeneration. Front Cell Dev Biol 7, 296 (2019).

80. Mitchell, D.M., Lovel, A.G. & Stenkamp, D.L. Dynamic changes in microglial and macrophage characteristics during degeneration and regeneration of the zebrafish retina. J Neuroinflammation 15, 163 (2018).

81. Conedera, F.M., Pousa, A.M.Q., Mercader, N., Tschopp, M. & Enzmann, V. Retinal microglia signaling affects Muller cell behavior in the zebrafish following laser injury induction. Glia 67, 1150–1166 (2019).

82. White, D.T. et al. Immunomodulation-accelerated neuronal regeneration following selective rod photoreceptor cell ablation in the zebrafish retina. Proc Natl Acad Sci U S A 114, E3719–E3728 (2017).

